# Identifying A- and P-site locations on ribosome-protected mRNA fragments using Integer Programming

**DOI:** 10.1101/490755

**Authors:** Nabeel Ahmed, Pietro Sormanni, Prajwal Ciryam, Michele Vendruscolo, Christopher M. Dobson, Edward P. O’Brien

**Author notes:** These authors contributed equally to this work (Nabeel Ahmed, Pietro Sormanni). Present address: Department of Neurology, Columbia University College of Physicians and Surgeons, New York, NY, USA.

## Abstract

Identifying the A- and P-site locations on ribosome-protected mRNA fragments from Ribo-Seq experiments is a fundamental step in the quantitative analysis of transcriptome-wide translation properties at the codon level. Many analyses of Ribo-Seq data have utilized heuristic approaches applied to a narrow range of fragment sizes to identify the A-site. In this study, we use Integer Programming to identify A-site by maximizing an objective function that reflects the fact that the ribosome’s A-site on ribosome-protected fragments must reside between the second and stop codons of an mRNA. This identifies the A-site location as a function of the fragment’s size and its 5□ end reading frame in Ribo-Seq data generated from *S. cerevisiae* and mouse embryonic stem cells. The correctness of the identified A-site locations is demonstrated by showing that this method, as compared to others, yields the largest ribosome density at established stalling sites. By providing greater accuracy and utilization of a wider range of fragment sizes, our approach increases the signal-to-noise ratio of underlying biological signals associated with translation elongation at the codon length scale.

## Introduction

Translation is a fundamental cellular process and an important step of gene expression resulting in the production of proteins in cells^1^. In the past decade the advent of Ribo-Seq (also known as Ribosome profiling), a high-throughput Next-Generation Sequencing method^2,3^, has enabled the transcriptome-wide study of translation. Ribo-Seq involves rapidly halting translation in cells through the use of antibiotics or flash freezing followed by cell lysis and then digestion of the lysate using an RNase enzyme^4^. The resulting pool of ribosome-protected mRNA fragments is then amplified and sequenced. The number and length of mRNA fragments that map to the coding sequences (CDSs) of transcripts is a function of the location and number of ribosomes that were sitting at a particular location on different copies of the same transcript. Where the ribosome’s A- and P-sites were located on a fragment during the digestion step is not known *a priori*, additional information and assumptions must be introduced to estimate their locations. Since translation occurs at the A- and P-sites, the identification of these sites is critical to address translation-related questions. If the A- and P-sites are not accurately identified, then systematic or random error can diminish the statistical power of any underlying biological signal that might exist. The identification of the A- and P-sites within ribosome footprints is therefore fundamental to quantitatively understanding translation at the codon length scale.

Because of the importance of this assignment problem, a number of methods for identifying the A- and P-sites have been created^2,5–13^. Many of these approaches utilize the biological fact that only the P-site is permitted to occupy the start codon during translation initiation and only the A-site is permitted to occupy the stop codon during termination. Using such approaches, the A-site location in *S. cerevisiae* Ribo-Seq datasets, for example, has been estimated to be 15 nt from the 5□ end of ribosome-protected mRNA fragments of size 28 nt^2,14^; 16 nt for fragment size 29 nt^14^; 15 nts from the 5□ end of fragments that are 30 nt in length^15^ and frame-specific offsets of 14 to 17 nts from the 5□ end for fragments between 28 and 30 nt in length^12,16^. The P-site location offset is 3 nt prior to the A-site. Similarly, in mouse embryonic stem cells (mESCs), such approaches have yielded specific offsets for different fragment lengths^11^.

Here, we utilize the fundamental biological fact that the A-site on ribosome-protected fragments must reside within the CDS of a gene under normal growth conditions and without any upstream open reading frames. We use this fact to create an objective function that, when maximized, identifies where the ribosome’s A- and P-sites are most likely to be located on a ribosome-protected mRNA fragment. We apply our method to *S. cerevisiae* and mESCs Ribo-Seq datasets and show that, compared to other methods, our approach has greater accuracy and statistical power in identifying A- and P-site locations and assigning read density.

## Methods

### Integer Programming Algorithm

In the analysis of Ribo-Seq data, mRNA fragments are initially aligned onto the reference transcriptome and their location is reported with respect to their 5 end. This means that one fragment will contribute one read that is reported on the genome coordinate to which the 5 end nucleotide of the fragment is aligned (Fig. 1A). In Ribo-Seq data, fragments of different lengths are observed that can arise from incomplete digestion of RNA and from the stochastic nature of mRNA cleavage by the RNase used in the experiment (Figs. 1C-D, Supplementary Fig. S1). A central challenge in quantitatively analyzing Ribo-Seq data is to identify from these Ribo-Seq reads where the A- and P-sites were located at the time of digestion. It is non-trivial to do this since incomplete digestion and stochastic cleavage can occur at both ends of the fragment. For example, mRNA digestion resulting in a fragment of size 29 nt can occur in different ways, two of which are illustrated in Fig. 1B. The quantity that we need to accurately estimate is the number of nucleotides that separate the codon in the A-site from the 5□ end of the fragment, which we refer to as the offset and denote Δ. Knowing Δ determines the position of the A-site as well as the P-site since the P-site will always be at Δ minus 3 nt.

**Figure 1:**
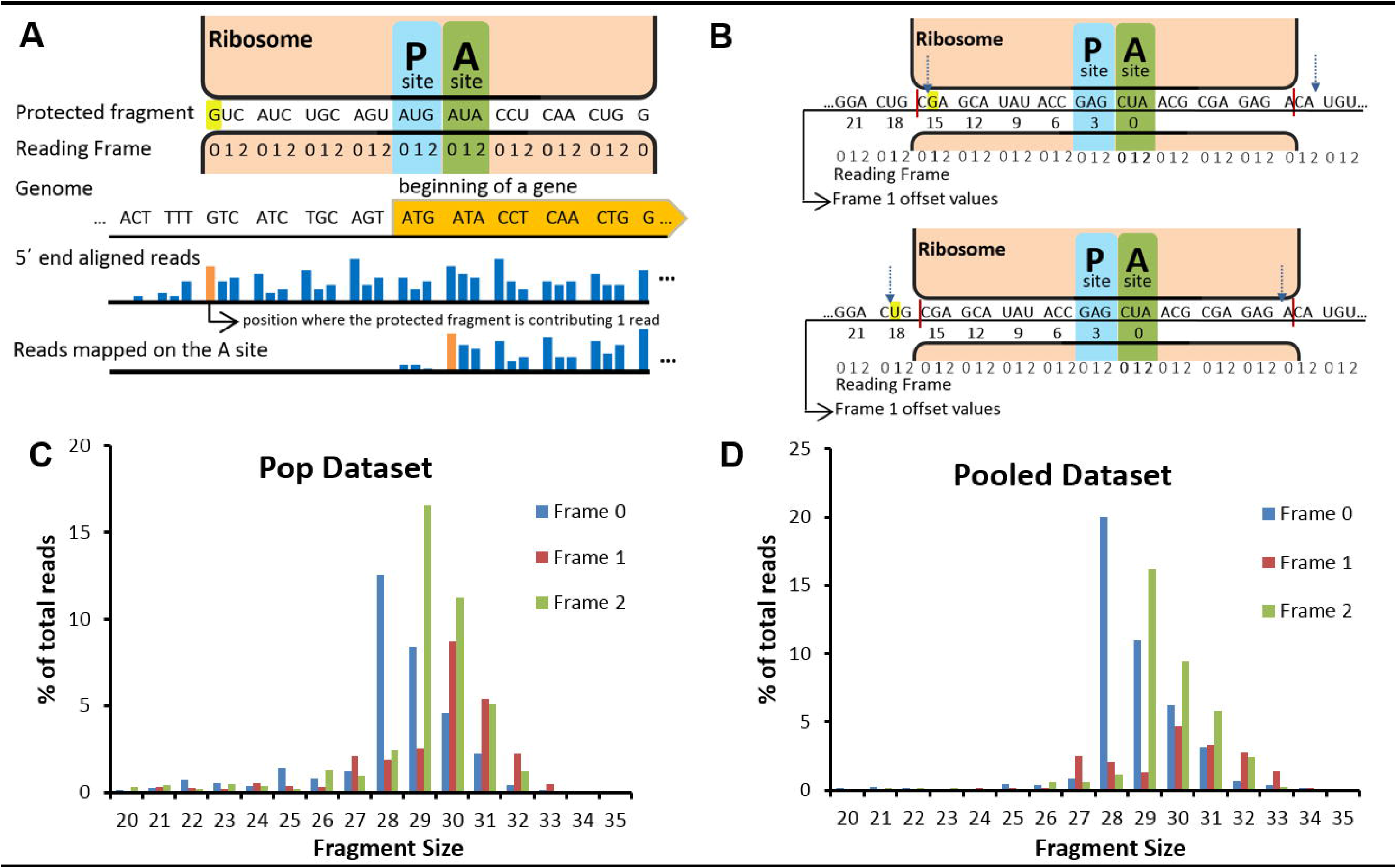
The A-site location can be defined as an offset from the 5□ end of ribosome-protected fragments. **(A)** A schematic representation of a ribosome beginning translation (top drawing) and of the offset between the Ribo-Seq reads mapped with respect to the 5 end of footprints and centered on the A-site (orange bar plots). The ribosome is shown protecting a 28 nt fragment with its 5 end in reading frame 0. The start codon of a gene can only occupy the P-site and hence the A-site was determined to be at an offset of 15 nt from the 5□ end for fragment size 28 which is in frame 0 (highlighted in yellow)^2^. The P-site and A-site within the fragment are indicated. The reads are then shifted from the 5□ end to the A-site by the offset value. **(B)** The boundaries of the 28 nt ribosome-protected footprint are indicated by red bars. Stochastic nuclease digestion can result in different fragment sizes. Two most probable variants of a 29 nt footprint with the 5□ end in frame 1 (highlighted in yellow) are shown with dashed arrows which can result in offsets of 15 nt (top) and 18 nt (bottom), respectively. (C-D) mRNA fragment size distribution for *S. cerevisiae* Ribo-Seq dataset from Pop and co-workers **(C)** and the Pooled dataset **(D)**

Our solution to this problem relies on the biological fact that for canonical transcripts with no upstream translation the A-site of actively translating ribosomes must be located between the second codon and stop codon of the CDS of a transcript^17^. Therefore, the optimal offset value Δ for fragments of a particular size (S) and reading frame (F) that map onto gene *i* is the one that maximizes the total number of reads *T*(Δ|*i,S,F*) between these codons. The size of an mRNA fragment *S* is measured in nucleotides, and the frame has values of 0, 1 or 2 and corresponds to the frame in which the 5 end nucleotide of the fragment is located. The 5 end frame is a result of RNase digestion and it is distinct from the reading frame of the ribosome that is typically translating in-frame (frame 0 of A-site). This concept can be expressed in terms of Integer Programming^18^, a mathematical optimization procedure in which an objective function is maximized subject to integer and linear restraints. With Δ as the integer variable to optimize, the objective function in this case is 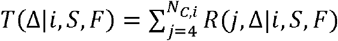, where *N_C,i_* is the number of nucleotides in the CDS of gene *i* and *R*(*j*, Δ|*i,S,F*) is the number of reads from fragments of size *S* and frame *F* mapped onto gene *i* whose 5□ end is at nucleotide position on the CDS after being shifted along the transcript by Δ nucleotides. The optimal Δ, denoted Δ′, for a given (*S, F*) for gene is determined as max{*T*(Δ|*t,S,F*)} subject to the constraints (*i*) that 0≤Δ≤S, and (*ii*) that the modulus of 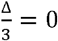. Constraint (*i*) enforces the requirement that the A-site is located between the first and last nucleotide of the fragment of size *S* nts. Constraint (*ii*) maintains the frame of the 5□-most nucleotide of the fragment as the Ribo-Seq reads are shifted by an amount Δ. We enforce Constraint (*ii*) because we are interested in the assignment of reads to the A-site at the resolution of a codon, not an individual nucleotide. If we did not enforce constraint (*ii*), our algorithm would simply yield equal *T*(Δ|*i, S, F*) scores for the two other values of Δ that would still map the reads on the A-site codon, but in the two frames where the 5 end was not in. Therefore, to simplify the determination of offsets we implemented constraint (*ii*). Thus, by maximizing *T*(Δ|*i, S, F*) for the CDS of each gene in a data set of *N_g_* genes, we will obtain a set of *N_g_* values of Δ’. From this distribution of Δ’ values, the A-site location corresponds to the most probable Δ’ value.

While identifying the Δ′ value for each gene in our data set, we also minimize the occurrence of false positives by ensuring that the highest score, *T*(Δ′|*i,S,F*), is significantly higher than the next highest score, *T*(Δ″|*i,S,F*), which occurs at a different offset Δ″. If the difference between the top two scores is less than the average number of reads per codon, we apply the following additional selection criteria. To choose between Δ′ and Δ″, we select the one that yields a number of reads at the start codon that is at least one-fifth less than the average number of reads at the second, third and fourth codons. We further require that the second codon have a greater number of reads than the third codon. The biological basis for these additional criteria are that the true offset (*i.e.*, the actual location of the A-site) cannot be located at the start codon, and that the number of reads at the second codon should be higher on average than the third codon due to contributions from the initiation step of translation, during which the ribosome is assembling on the mRNA with the start codon in the P-site. In the Results section, we demonstrate that the results from our method are largely robust to changes in these thresholds.

### Ribo-Seq datasets

#### S. cerevisiae

Published Ribo-Seq data from *S. cerevisiae* were obtained from GSM1557447 used in the study of Pop and co-workers^19^. The raw reads were pre-processed according to the method stated in the original study. Raw fastq files were downloaded and preprocessed using Fastx-toolkit (v0.013) (http://hannonlab.cshl.edu/fastx_toolkit/index.html) as stated in the methods of the original study. The adapter sequence CTGTAGGCACCATCAAT was stripped using FastQ clipper and low-quality reads were filtered by FastQ quality filter. The processed reads were aligned first to the ribosomal RNA sequences using Bowtie 2 (v2.2.3)^20^. The reads which did not align to the ribosomal sequences were then aligned to the *Saccharomyces cerevisiae* assembly R64-2-1 (UCSC: sacCer3) using Tophat (v2.0.13)^21^ with up to two mismatches allowed. Gene annotations were obtained from Saccharomyces Genome Database (http://www.yeastgenome.org/) on May 4, 2016 for 6,572 protein-coding genes. Reads were assigned to the nucleotide positions according to the 5□ end.

The pooled Ribo-Seq dataset was formed by combining reads from all replicates *of S. cerevisiae* Ribo-Seq data published in studies in which cycloheximide (CHX) was not used to induce translation arrest^14–16,19,22–28^. It has been demonstrated that CHX pre-treatment leads to distortion of ribosome profiles due to ribosome slippage even after CHX treatment^12,22^. The distorted ribosome profiles can spill across the CDS boundaries thus limiting the application of Integer Programming algorithm. Hence, our analysis only used those datasets without CHX pre-treatment. The list of all the utilized datasets is reported in Supplementary Table S1. The raw reads from each study were processed according to the reported method in the original study. If the method is not reported in the original study, we used cutadapt (v1.14)^29^ to pre-process the raw reads. The alignment and assignment of reads to gene transcripts was done as above for the Pop dataset^19^.

#### Mouse embryonic stem cells

The “no drug” sample for mouse embryonic stem cells (mESCs) measured by Ingolia and co-workers^11^ was utilized in this study. Since CHX treatment has been shown to artificially alter ribosome profiles in *S. cerevisiae*, we believed it prudent to not use mESC samples pretreated with CHX. To increase the coverage we pooled reads from another untreated Ribo-Seq sample of mESCs published in the study of Hurt and co-workers^30^. The linker sequence CTGTAGGCACCATCAATTCGTATGCCGTCTTCTGCTTGAA for Ingolia’s dataset and the poly-A adapter sequence for Hurt’s dataset were trimmed using cutadapt (v1.14)^29^. The trimmed reads were first aligned to ribosomal RNA sequences using Bowtie2 (v2.2.3)^20^ and the filtered reads were subsequently aligned to mm10 reference transcriptome consisting of 21,185 genes obtained from UCSC knownGene database using Tophat (v2.0.13)^21^ with up to two mismatches allowed. For a gene with multiple isoforms, only the isoform with the longest CDS was included in the reference transcriptome. For transcripts with no information on the 5□ UTR region, we included 40 nt of genomic sequence upstream from the start codon for successful alignment of reads around start codon and effective application of Integer Programming algorithm. Translation initiation site data was obtained from Table S3 of study of Ingolia and co-workers^11^. We selected genes that have only one translation initiation site coding for only a canonical CDS product. From these genes, only genes containing a single isoform were selected, resulting in 430 genes in our final dataset.

#### Escherichia coli

Wild-type Ribo-Seq data for *E.coli* were obtained from studies of Li and co-workers (2012)^31^, Li and co-workers (2014)^32^ and Woolstenhulme and co-workers^33^. The accession numbers of the samples used are provided in Supplementary Table S1. The respective linker sequences in each sample were trimmed using cutadapt (v1.14)^29^. Reads were initially aligned to ribosomal RNA sequences using Bowtie2 (v2.2.3)^20^ and the rest of reads aligned to the *E.coli* reference genome build NC_000913.3 using Tophat (v2.0.13)^21^ with up to two mismatches allowed. Gene annotations were obtained for 4314 genes from RefSeq database corresponding to NC_000913.3.

### Gene selection, analyses and statistical tests

#### Selection of genes

To obtain good sampling statistics, we selected for analysis only those genes that have on average greater than 1 read per codon per fragment length per reading frame. This means that different sets of genes can be used in the Integer Programming algorithm depending on the fragment length and frame under scrutiny. The average number of reads per codon was calculated on the CDS region of the gene and an additional upstream region corresponding to the size of the fragment length being considered. Genes in which more than 1% of the total number of mapped reads, for a given *S* and F, mapped to multiple locations across the genome were discarded from further analysis.

#### Identifying unique offsets

We defined the most probable offset Δ to have a unique, unambiguously identified A-site if at least 70% of genes in the dataset had an offset equal to Δ’, and further require that there be at least 10 genes in the dataset. Otherwise, the A-site location is defined as ambiguous for the fragment size and frame under scrutiny. In the Results section, we show the A-site location is largely robust to moderate variation in this 70% threshold.

#### High coverage test

To test for the effect of depth of coverage on the A-site location we increased the average number of reads per codon required for a gene to be included in the analyzed dataset from 1 to values up to 50. Three requirements have to be met for an ambiguous offset to be identified as unique as coverage is increased. As before, 70% of the genes had to have the most probable offset with at least 10 genes in the dataset. In addition, there must to be a statistically significant increasing trend in the most probable offset with increasing coverage. This requirement prevents fluctuations above 70% due to statistical error as being counted as a unique offset. This trend is calculated using Linear Regression Analysis.

#### Test using Artificial Ribo-Seq data

To construct artificial ribosome occupancies, we used Gillespie’s algorithm^34^ to simulate translation across *S. cerevisiae* mRNA transcripts. During the simulations, we saved snapshots every X steps recording the A-site codon location and creating a histogram of ribosome occupancies across the transcript. To be consistent with the sampling statistics of the experimental Pooled *S. cerevisiae* data, we carried out our analysis on the same 4,487 transcripts that met our filtering criteria for (*S, F*) = (28,0), and normalized our simulated ribosome occupancies such that they sum up to the total number of reads mapped to that transcript in the experimental data. We then created different fragment size and 5 end reading frame distributions (Supplementary Fig. S3A, B). Specifically, since the reads are counts, we use Poisson statistics by treating each (*S, F*) as an event in the order: (20,0), (20,1),(20,2), …, (35,0), (35,1) and (35,2). Six Poisson distributions of different variances (1 = 4,8,16,24,48,80) were generated. The distributions were shifted such that the mode of the distribution was at (*S, F*) = (28,0), which is typically found in experiments, with probabilities summing up to 1 between (20,0) and (35,2). Two additional read length distributions were also considered with modes at (*S, F*) = (24,0) and (*S, F*) = (32,0) withl = 8. Four different sets of offset tables were used as an input to generate the artificial Ribo-Seq reads from the simulated ribosome occupancies for each of these distributions. These four offset sets are i) a constant offset of 15 nucleotides for all (*S, F*)s, ii) a constant offset of 18 for all (*S, F*)s iii) a constant offset of 12 for *S* = 20,21, …, 26,27 and constant offset of 18 for *S* = 28,29, …, 34,35 iv) the “top offset” values for (*S, F*) combinations identified using our algorithm in the experimental Pooled *S. cerevisiae* data (*i.e.*, the offset values of Table 1). These input offset tables were compared to the ‘output’ offset table generated by applying the IP algorithm on the artificial Ribo-Seq data to test the correctness of our method.

**Table 1:** A-site locations (nucleotide offsets from 5 end) determined by applying the Integer Programming algorithm to the Pooled dataset in *S. cerevisiae* are shown as a function of fragment size and frame. The top two offset values are listed for those *S* and F combinations in which the A-site location could not be uniquely determined. For unique offsets, the most-probable offset value is listed.

| Fragment Size | Frame 0 | Frame 1 | Frame 2 |
| --- | --- | --- | --- |
| 24 | 15 | 15/12 | 18/12 |
| 25 | 15 | 12/15 | 18 |
| 26 | 15/12 | 18/15 | 18/15 |
| 27 | 15 | 15 | 18 |
| 28 | 15 | 15 | 18 |
| 29 | 15 | 15/18 | 18 |
| 30 | 15 | 18 | 18 |
| 31 | 15 | 18 | 18 |
| 32 | 18/15 | 18 | 18 |
| 33 | 18 | 18 | 18 |
| 34 | 18 | 18 | 18/21 |

#### Statistical significance of PPX and XPP motifs

To test if the normalized read density distribution of a PPX or XPP motif is not due to random chance, we calculated the P-value using a permutation test^35^. For the total number of instances of a PPX/XPP motif, we randomly selected an equal number of instances of any other three-residue motif and determined the median normalized read density at the third codon position of the motif, thereby creating a random distribution. We repeated this procedure 10,000 times and calculated the fraction of iterations that had a median density equal to or greater than the one observed for that PPX/XPP motif. This fraction is equal to the P-value. The instances of PPX and XPP motifs are identified from those transcripts that have at least 50% of codon positions with 1 read or more.

#### Comparison with other A-site mapping methods

We compared the performance of Integer Programming algorithm with other methods by calculating the difference in normalized read density between the Integer Programming A-site value and the compared method’s A-site value at the third codon of PPG and PPE motifs, which are associated with ribosome pausing in *S. cerevisiae* and mESCs respectively.

In *S. cerevisiae*, A-site ribosome profiles were obtained for Integer Programming method by applying the offsets listed in Table 1 for fragment sizes 24 to 34 nt. For methods used by Martens and co-workers and Hussmann and co-workers^12^ specifically in *S. cerevisiae*, A-site profiles were obtained by applying the offsets for specific fragment sizes as stated in the Methods sections of those studies. We included a constant heuristic offset of 15 nt which has been used in several studies of *S. cerevisiae* Ribo-Seq data^2,36–38^. The constant offset of 15 nt has been applied to a wide range of fragment lengths across studies including 22-32 nt^2^, 27-30 nt^36^, 28 nt^37^, 27-34 nt^38^. To be conservative, we apply a constant offset of 15 nt to fragments between 27 and 30 nt only. Similarly, we also include a method where a constant offset of 18 nt is applied to fragments between 27 and 30 nt to compare to the performance of the Integer Programming method.

For mESCs, Ingolia and co-workers^11^ implemented length specific offsets of 15, 16 and 17 nts from the 5□ end, respectively, for fragments of size 29-30 nt, 31-33 nt and 34-35 nt. Several studies have also implemented a constant offset of 15 for range of fragment sizes 25-35 nt^39,40^. Similar to *S. cerevisiae*, we also implement a constant offset of 18 nt to fragment size range of 25-35 nt.

Few general methods have been proposed to determine A-site locations in any organism. We implemented the methods Plastid^7^, RiboProfiling^8^ and riboWaltz^9^ which are publicly available as R packages. The A-site offset tables generated using these methods for our analyzed datasets in *S. cerevisiae* and mESCs are presented in Supplementary Table S9. To determine the A-site profiles using the ‘ribodeblur’ method created by Wang and co-workers^6^, we ran the source code available in GitHub (https://github.com/Kingsford-Group/ribodeblur-analysis/releases/tag/v0.1) on our datasets and added a custom Python script to generate the ‘deblurred’ A-site profiles. For Rpbp^41^, the publicly available software was downloaded and run locally to obtain the A-site offsets. We also applied the center-weighted method as described by Becker and co-workers^42^; for reads greater than 23 nt, we trim 11 nt from both ends of the fragment and distribute the read equally among the remaining nucleotides. For scikit-ribo method^10^, the source code was downloaded and was successfully run for *S. cerevisiae* datasets to obtain the A-site profiles. Scikit-ribo could not be run on mouse ESC data as the current available version of the source code contains bugs resulting in inaccurate annotation assignments for higher eukaryotic genomes.

Instances of PPG motifs (in *S. cerevisiae*) and PPE motifs (in mESCs) used for analysis are selected from genes in which at least 90% of codon positions have at least 1 read in their 5□ aligned ribosome profiles in the CDS region and an upstream region of 18 nt. An instance of a motif is included for analysis only if its ribosome density is greater than 1.5 of average ribosome density at the third codon position in the A-site profile of any compared methods. We use the Wilcoxon signed rank test to determine if there is a statistically significant difference between the normalized read density at the third codon of motif instances obtained by Integer Programming and other methods.

## Results

### Illustrating the Integer Programming optimization procedure

To illustrate this Integer Programming algorithm in action we provide an example using the hypothetical mRNA shown in Figure 2. The algorithm is as follows: First, for gene *i*, consider *RP*(*j*, Δ = 0|*i, S, F*)composed of those fragments of size *S* (= [20, 21, …, 35]nt) and whose 5 end has been aligned to reading frame *F* (=0,1 or 2). Second, for this ribosome profile, determine the Δ that maximizes *T*(Δ|*i, S, F*). Do this by starting from the 5□-end-aligned ribosome profile (Δ=0) and shift it three nucleotides at a time (*i.e*., obey Constraint 2 described in Methods) towards the 3 end of the transcript such that Δ = 0,3,6,9, …, ≤ *S*. At each value of Δ, calculate *T*(Δ|*i,S,F*) and record its value. Third, after all Δ values have been tested, the Δ that maximizes is denoted Δ′, which is the putative location of the A-site relative to 5□ end of fragments of size *S* and frame *F* for gene *i*. Check if the secondary-selection criteria are required and apply them when the scores for the top two offsets differ by less than the average number of reads per codon in the mRNA. Finally, repeat these steps for every fragment size between 20-35 nts in length and every reading frame. Thus, for one gene, this procedure yields 48 (=16×3) independent values for Δ’, one for each fragment size and frame combination.

**Figure 2:**
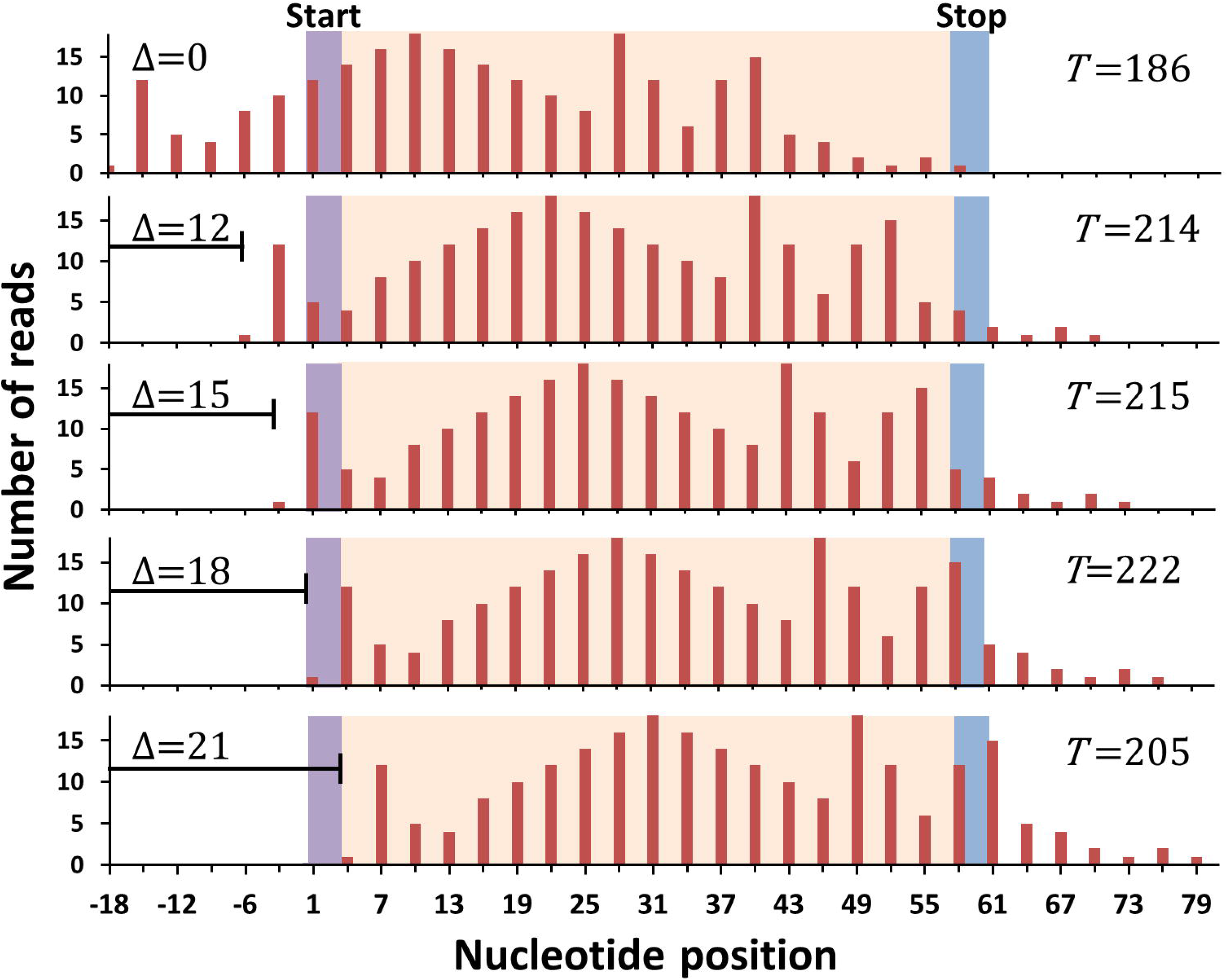
An illustration of the application of the Integer Programming algorithm to a Ribosome profile. For a hypothetical transcript that is 60 nt in length the first panel shows the ribosome profile originating from reads assigned to the 5□ end of fragments of size 33 in frame 0. The start and the stop codon are indicated while the rest of the CDS region is colored light peach. The algorithm shifts this ribosome profile by 3 nt and calculates the objective function *T*(Δ|*i,S,F*). The extent of the shift is the offset Δ. Values of *T*(Δ|*i,S,F*) for Δ= 12,15,18,21 nts are indicated. In this example, the average number of reads per codon is 7.85. The difference between the top two offsets, 18 (*T*=222) and 15 (*T*=215), is less than the average. Hence, we check the secondary criteria (Methods). Offset 18 meets the criteria that the number of reads in the start codon is less than one-fifth of the average of reads in second, third and fourth codons and also that number of reads in the second codon is greater than reads in third codon. Hence, Δ=18 nt is the optimal offset for this transcript.

The fragment-size and frame distributions of ribosome-protected fragments (Figs. 1C, D) in *S. cerevisiae* are not gene dependent (Supplementary Fig. S2), and therefore, neither should be the offset values. Thus, the location of the A-site, relative to the 5□ end of a fragment of size *S* and frame *F*, corresponds to the most probable value of the offset across all the genes in the dataset.

### A-site locations in *S. cerevisiae* Ribo-Seq data are fragment size and frame dependent

We first applied the Integer Programming method to Ribo-Seq data from *S. cerevisiae* published by Pop and co-workers^19^. For each combination of *S* and *F* we first identified those genes that have at least 1 read per codon on average in their corresponding ribosome profile. The number of genes meeting this criterion is reported in Supplementary Table S2. We then applied the Integer Programming method to this subset of genes. The resulting distributions of Δ values are shown in Fig. 3A for different combinations of fragment length and frame. We only show results for fragment sizes between 27 and 33 nt because greater than 90% of reads map to this range (Fig. 1C). The most probable offset value for all fragment sizes between 20 to 35 nt is reported as an offset table (Supplementary Table S4).

**Figure 3:**
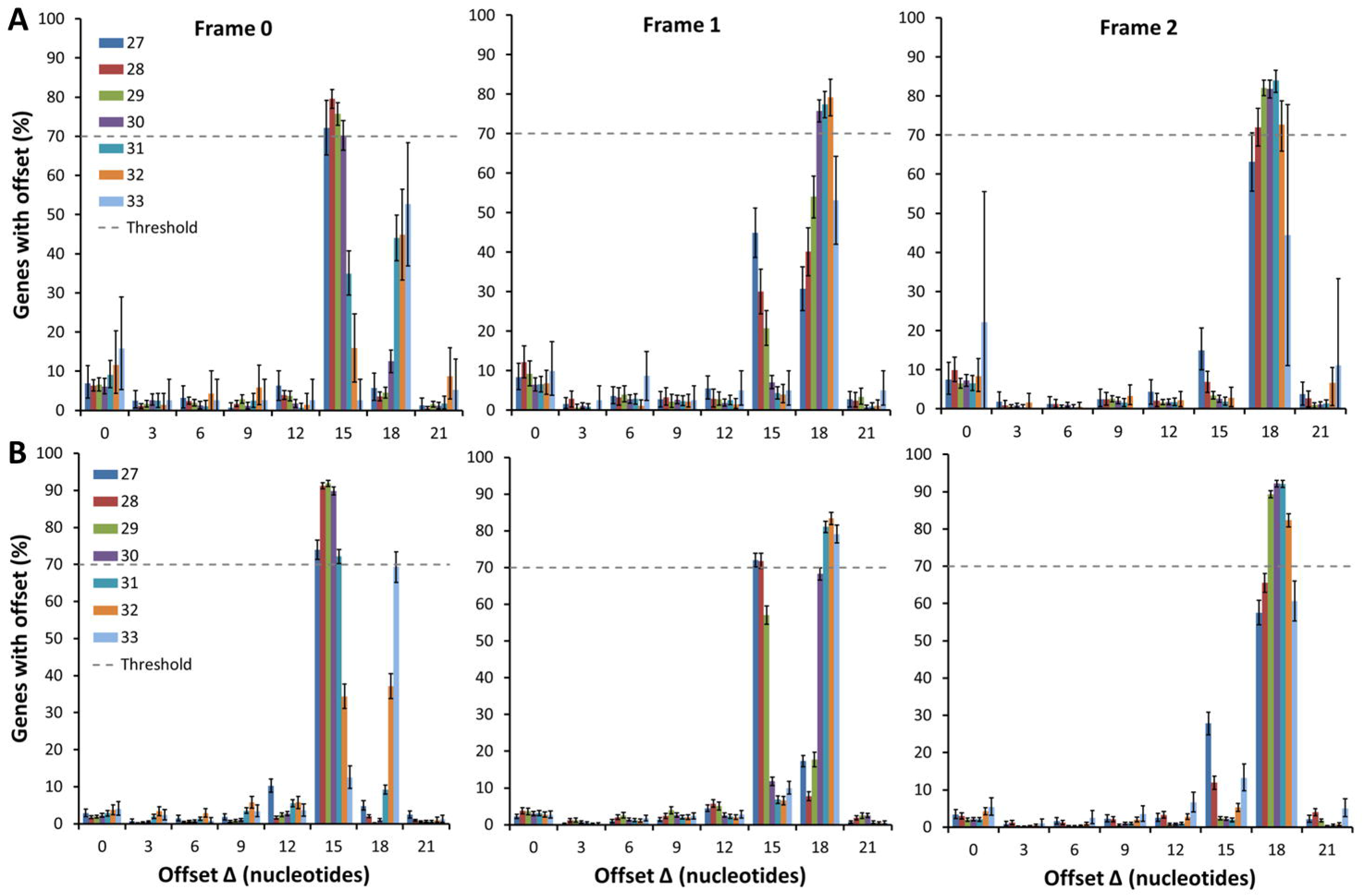
Distribution of offset values from the Integer Programming algorithm applied to transcripts from *S. cerevisiae*. The data plotted in **(A)** are from the Pop dataset, and **(B)** the Pooled dataset. The distributions are plotted as a function of the offset value and for fragment sizes of 27 to 33 nt, are shown, from left to right, for frames 0, 1 and 2. For a given fragment size and frame, the A-site location is at the most probable Δ value in the distribution, provided the offset occurs for more than 70% of the genes (dashed lines in panels). Error bars represent 95% Confidence intervals calculated using Bootstrapping. Sample sizes are reported in Supplementary Table S2.

We see that the optimal Δ value - that is, the A-site location - changes for different combinations of *S* and *F*, with the most probable values either at 15 or 18 nt. Thus, the location of the A-site depends on *S* and *F*. In most cases, there is one dominant peak for a given pair of *S* and *F* values. For example, for fragments of size 27 through 30 nt in frame 0, greater than 70% of their per-gene optimized Δ values are 15 nt from the 5□ end of these fragments. Similar results are found for other combinations such as sizes 30, 31 and 32 nt in frame 1 and 28 through 32 nt in frame 2, where optimized Δ values are 18 nt. Thus, across the transcriptome, the A-site codon position on these fragments is uniquely identified.

There are, however, *S* and *F* combinations that have ambiguous A-site locations based on these distributions. For example, for fragments of size 27 nt in frame 1, 47% of the gene-optimized Δ values are at 15 nt while 30% are at 18 nt. Similar results are observed for fragments 28 and 29 nt in frame 1, and 31 and 32 nt in frame 0. Thus, for these *S* and F combinations there is a similar probability of the A-site being located at one codon or another, and therefore we cannot uniquely identify the A-site’s location.

### Higher coverage leads to more unique offsets

We hypothesized that ambiguity in identifying the A-site for particular *S* and *F* combinations may be due to low coverage (*i.e.*, sampling poor statistics). To test this hypothesis we pooled the reads from different published Ribo-Seq datasets into a single dataset with consequently higher coverage and more genes that meet our selection criteria (Supplementary Table S2). Application of our method to this Pooled dataset gives unique offsets for more *S* and *F* combinations compared to the original Pop dataset (Fig. 3B and Supplementary Table S4), supporting our hypothesis. For example, for fragments of size 27 and frame 1, now we have the unique offset of 15 nt with 72% of gene-optimized Δ values at 15 nt (Fig. 3B). However, we still see the ambiguity present for certain (*S, F*) combinations.

We employed an additional strategy to increase coverage by restricting our analysis to genes with greater and greater average reads per codon. If the hypothesis is correct, then we should see a statistically significant trend of an increase in the most probable Δ value with increasing read depth. We applied this analysis to the Pooled dataset and find that some initially ambiguous *S* and *F* combinations become unambiguous as coverage increases. For example, at an average of 1 read per codon, (*S,F*) combinations of (25, 0), (27, 2), and (30, 1) are ambiguous as they fall below our 70% threshold. However, we see a statistically significant trend (*slope* = 0.5, *p* = 3.94 × 10^−6^) for fragments of that the 15 nt offset becomes more probable upon increasing the coverage, eventually crossing the 70% threshold (Fig. 4A). Similarly, for (27,2) (*slope* = 0.58, *p* = 5.77 × 10^−5^) and (30,1) (*slope* = 0.25, *p* = 0.009) there is a trend towards an offset of 18 nt, with more than 70% of genes having this offset at the highest coverage (Figs. 4B, C). Hence, for these fragments, increasing coverage uniquely identifies and hence the A-site location. For a few combinations of (*S, F*), like (32,0), the ambiguity is not resolved even upon very high coverage (Fig. 4D), which we speculate may be due to inherent features of nuclease digestion being equally likely for more than one offset.

**Figure 4:**
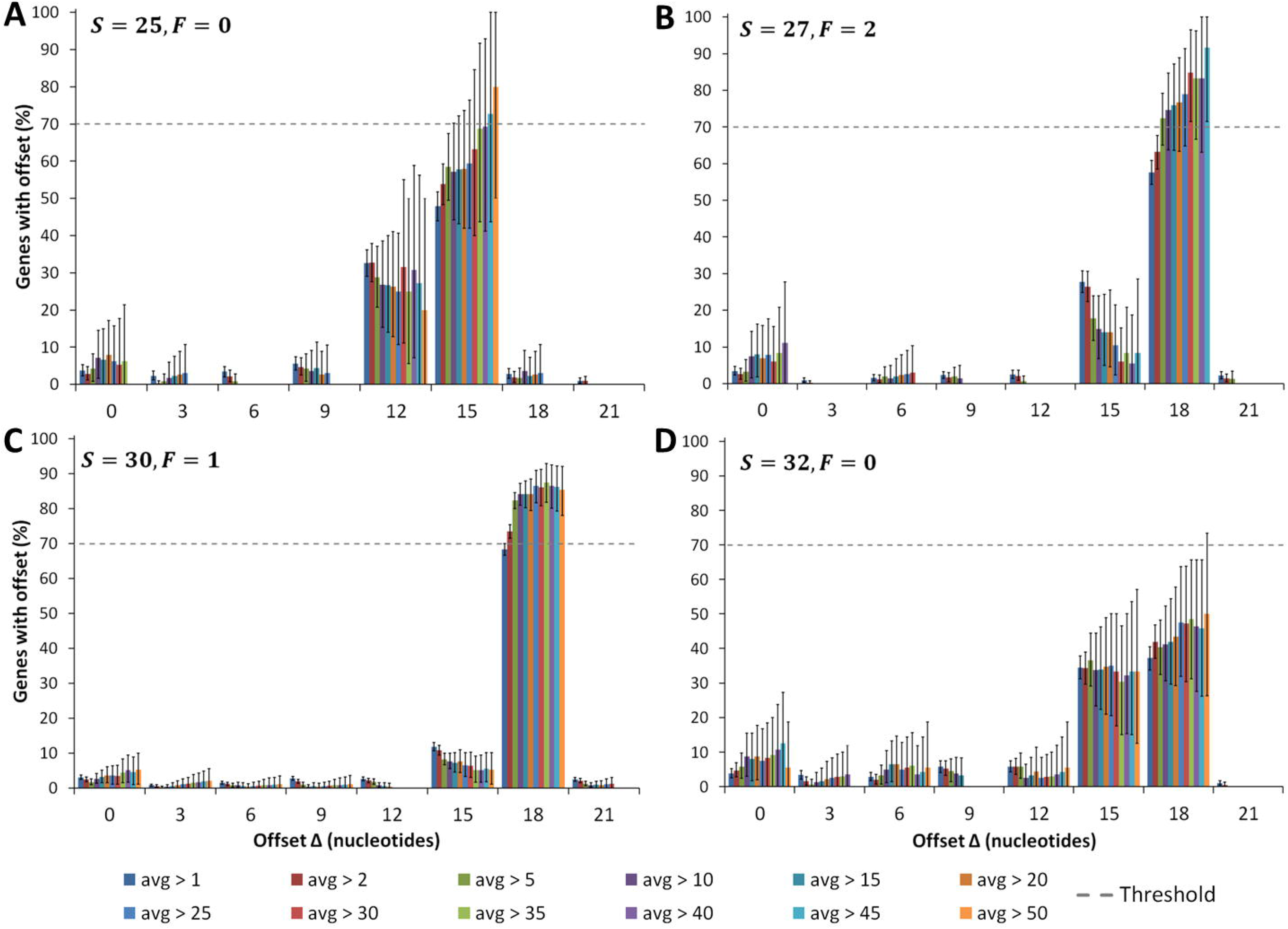
Increasing coverage identifies A-site locations for *S* and *F* combinations that were initially ambiguous. Plotted is the percentage of transcripts with a particular Δ value for different *S* and F combinations from the Pooled dataset of *S. cerevisiae*. In each panel, multiple distributions are plotted corresponding to transcripts with increasing coverage, indicated by the legend at the bottom. For example, the distributions in blue and red arise from transcripts with, respectively, at least 1 or 2 reads per codon on average. We observe the A-site location tends towards 15 nt for **(A)** and towards 18 nt for **(B)**, and **(C)**. For **(D)**, there is no trend even at higher coverage. Note that for *S* = 27, *F* = 2 (panel B), there are less than 10 genes with an average greater than 50 reads per codon and hence we do not include the data point beyond average greater than 45 reads per codon (see Methods). Error bars represent 95% Confidence intervals calculated using Bootstrapping.

Thus, high enough coverage yields the optimal offset table represented in Table 1, where the offset is the most probable location of the A-site relative to the 5 end of the mRNA fragments generated in *S. cerevisiae*.

### Consistency across different datasets

Ribo-Seq data is sensitive to experimental protocols that can introduce biases in the digestion and ligation of ribosome-protected fragments. Pooling datasets together offers the advantage of higher coverage but it may mask the biases specific to an individual dataset. To determine whether our unique offsets (Table 1) are consistent with results from individual data sets we applied the Integer Programming algorithm to each individual dataset. Most of these datasets have low coverage resulting in fewer genes meeting our filtering criteria (Supplementary File S1). For each unique offset in Table 1, we classify it as consistent with an individual data set provided that the most probable offset from the individual dataset (even if it does not reach the 70% threshold due to limitations in the depth of coverage) is the same as in Table 1. We find that the vast majority of unique offsets (18 out of 20) in Table 1 are consistent across 75% or more of the individual datasets (statistics reported in Supplementary Table S5). Just two (*S,F*) combinations show frequent inconsistencies. (*S,F*) combinations (27,1) and (27,2) are inconsistent in 33% or more of the individual datasets (Supplementary Table S5). This suggests that researchers who wish to minimize false positives should discard these (*S, F*) combinations when creating A-site ribosome profiles.

### Robustness of the offset table to threshold variation

The Integer Programming algorithm utilizes two thresholds to identify unique offsets. One is that 70% of genes exhibit the most probable offset, the other, designed to minimize false positives arising due to sampling noise in the Ribo-Seq data, is that the reads in the first codon be less than one-fifth of the average reads in the second, third and fourth codon. While there are good reasons to introduce these threshold criteria, the exact values of these thresholds are arbitrary. Therefore, we tested whether varying these thresholds changes the results reported in Table 1. We varied the first threshold to 60% and 80%, and recomputed the offset table. We report whether the unique offset changed by listing an ‘R’ or ‘S’ (for robust and sensitive, respectively) alongside the reported offset in Supplementary Table S5. We find that two-thirds of the unique (*S,F*) combinations do not change (Supplementary Table S5). (*S,F*) combinations (25, 0), (25, 2), (27, 0), (27, 1), (28, 1), (31, 0), (33, 0) and (33, 2) become ambiguous when we increased the threshold to 80%.

We varied the second, aforementioned threshold from one-fifth up to one and down to one-tenth, and we find that all unique (*S,F*) combinations except (25,2), (33,0), (33,2) and (34,1) remain unchanged (reported as ‘R’ in Supplementary Table S5). Thus, in summary, in the vast majority of cases, the unique offsets reported in Table 1 depend very little on specific values of these thresholds.

### Testing the Integer Programming algorithm against artificial Ribo-Seq data

To test the correctness and robustness of our approach we generated a dataset of simulated ribosome occupancies across 4,487 *S. cerevisiae* transcripts and asked whether our method could accurately determine the A-site locations. Artificial Ribo-Seq reads were generated from these occupancies assuming a Poissonian distribution in their (*S, F*) values using random footprint lengths similar to that found in experiments (see Methods and Supplementary Fig. S3A, B). We investigated the ability of our method to correctly determine the true A-site locations for four different sets of pre-defined offset values (see Methods). The Integer Programming algorithm was then applied to the resulting artificial Ribo-Seq data. We find the offset table generated from the algorithm reproduces the input offsets used (Supplementary Fig. S3C and Supplementary Table S6). This procedure was repeated for different read length distributions as well as with different input offsets and we find that the offset tables generated by our algorithm reproduce the input offset tables in greater than 93% of all (*S, F*) combinations (Supplementary Fig. S3B, C and Supplementary File S2). The method identifies a small number of ambiguous offsets due to the low read coverage at the tails of the distributions. A finding that emphasizes further the importance of read coverage as a critical factor in accurately identifying the A-site.

### A-site offsets in mouse embryonic stem cells

The biological fact that A-site of a ribosome resides only between the second and stop codon is not limited to *S. cerevisiae* and hence the Integer Programming algorithm should be applicable to Ribo-Seq data from any organism. Therefore, we applied our method to a Pooled Ribo-Seq dataset of mouse embryonic stem cells (mESCs). The resulting A-site offset table exhibited ambiguous offsets at all but three (*S,F*) combinations (Supplementary Table S7). In mESCs there is widespread translation elongation that occurs beyond the boundaries of annotated CDS regions in upstream open reading frames (uORFs)^39^. Enrichment of ribosome-protected fragments from these translating uORFs can make it difficult for our algorithm to find unique offsets because they can contribute reads around the start codon of canonical annotated CDSs. Therefore, we hypothesized that if we apply our algorithm to only those transcripts devoid of uORFs and possessing a single initiation site then our algorithm should identify more unique offsets. Ingolia and co-workers^11^ have experimentally identified for well-translated mESCs transcripts its number of initiation sites and whether uORFs are present using translation-initiation inhibiting drug Harringtonine. Therefore, we selected those genes that have only one translation initiation site near the annotated start codon and further restricted our analysis to transcripts with a single isoform, as multiple isoforms can have different termination sites.

Application of Integer Programming algorithm to this set of genes increases the number of unique offsets from 3 to 13 (*S,F*) combinations (Supplementary Table S8). Applying the same robustness and consistency tests as we did in *S. cerevisiae* reveals that 77% of the unique offsets are robust to threshold variation, and a similar percentage is consistent across both individual datasets used to create the Pooled data (Supplementary Table S8). Thus, the unique offsets we report for mESCs are robust and consistent in the vast majority of datasets. This result also indicates that successful identification of A-site locations requires analysing only those transcripts that do not contain uORFs.

### Integer Programming does not yield unique offsets for *E.coli*

As a further test of how widely we can apply our algorithm, we applied it to a Pooled Ribo-Seq data from the prokaryotic organism *E. coli*. The number of genes meeting our filtering criteria is reported in Supplementary Table S3. MNase, the nuclease used in the *E. coli* Ribo-Seq protocol, digests mRNA in a biased manner - favoring digestion from the 5□ end over the 3□ end ^33,43^. Therefore, as done in other studies^33,43,44^, we applied our algorithm such that we identified the A-site location as the offset from the 3□ end instead of the 5□ end. Polycistronic mRNAs (*i.e.*, transcripts containing multiple CDSs) can cause problems for our algorithm due to closely spaced reads at boundaries of contiguous CDS being scored for different offsets in both the CDSs. To avoid inaccurate results, we restrict our analysis to the 1,915 monocistronic transcripts that do not have any other transcript within 40 nt upstream or downstream of the CDS. Based on our experience in the analysis of mESCs dataset, we filter out transcripts with multiple translation initiation sites as well as transcripts whose annotated initiation sites have been disputed. Nakahigashi and co-workers^45^ have used tetracycline as translation inhibitor to identify 92 transcripts in *E.coli* with different initiation sites from the reference annotation and we exclude these transcripts from our analysis. However, for this high coverage pooled dataset, we find ambiguous offsets for all (*S,F*) combinations (Supplementary Table S7). A meta-gene analysis of normalized ribosome density in the CDS and 30 nt region upstream and downstream reveal signatures of translation beyond the boundaries of the CDS (Supplementary Fig. S4), especially a higher than average enrichment of reads a few nucleotides before the start codon. We speculate that the base-pairing of the Shine-Dalgarno (SD) sequence with the complementary anti-SD sequence in 16S rRNA^46^ protects these few nucleotides before the start codon from ribonuclease digestion and hence results in an enrichment of Ribo-Seq reads. Since these “pseudo” ribosome-protected fragments cannot be differentiated from actual ribosome-protected fragments containing a codon with the ribosome’s A-site on it, our algorithm is limited in its application for this data.

### Reproducing known PPX and XPP motifs that lead to translational slowdown

In *S. cerevisiae*^47^ and *E. coli*^33,48^ certain PPX and XPP polypeptide motifs (in which X corresponds any one of the 20 amino acids) can stall ribosomes when the third residue is in the A-site. Elongation factors eIF5A (in *S. cerevisiae*) and EF-P (in *E. coli*) help relieve the stalling induced by some motifs but not others^47^. Even in mESCs, Ingolia and co-workers^11^ detected PPD and PPE as strong pausing motifs. Therefore, we examined whether our approach can reproduce the known stalling motifs. We did this by calculating the normalized read density at the different occurrences of a PPX and XPP motif.

In *S. cerevisiae*, we observed large ribosome densities at PPG, PPD, PPE and PPN (Fig. 5A), all of which were classified as strong stallers in *S. cerevisiae*^47^ and also in *E. coli*^8^. In contrast, there is no stalling, on average, at PPP, consistent with other studies^47^. This is most likely due to the action of eIF5A. For the XPP motifs, the strongest stalling was observed for GPP and DPP motifs, which are consistent with the results in *S. cerevisiae* and in *E. coii* (Fig. 5B). In mESCs, we see the strongest stalling at PPE and PPD, reproducing the results of Ingolia and co-workers^11^ (Supplementary Fig. S5A). For XPP motifs, we observed very weak stalling only for DPP (Supplementary Fig. S5B). Thus, our approach to map the A-site on ribosome footprints enables the accurate detection of established translation pausing at particular PPX and XPP nascent polypeptide motifs.

**Figure 5:**
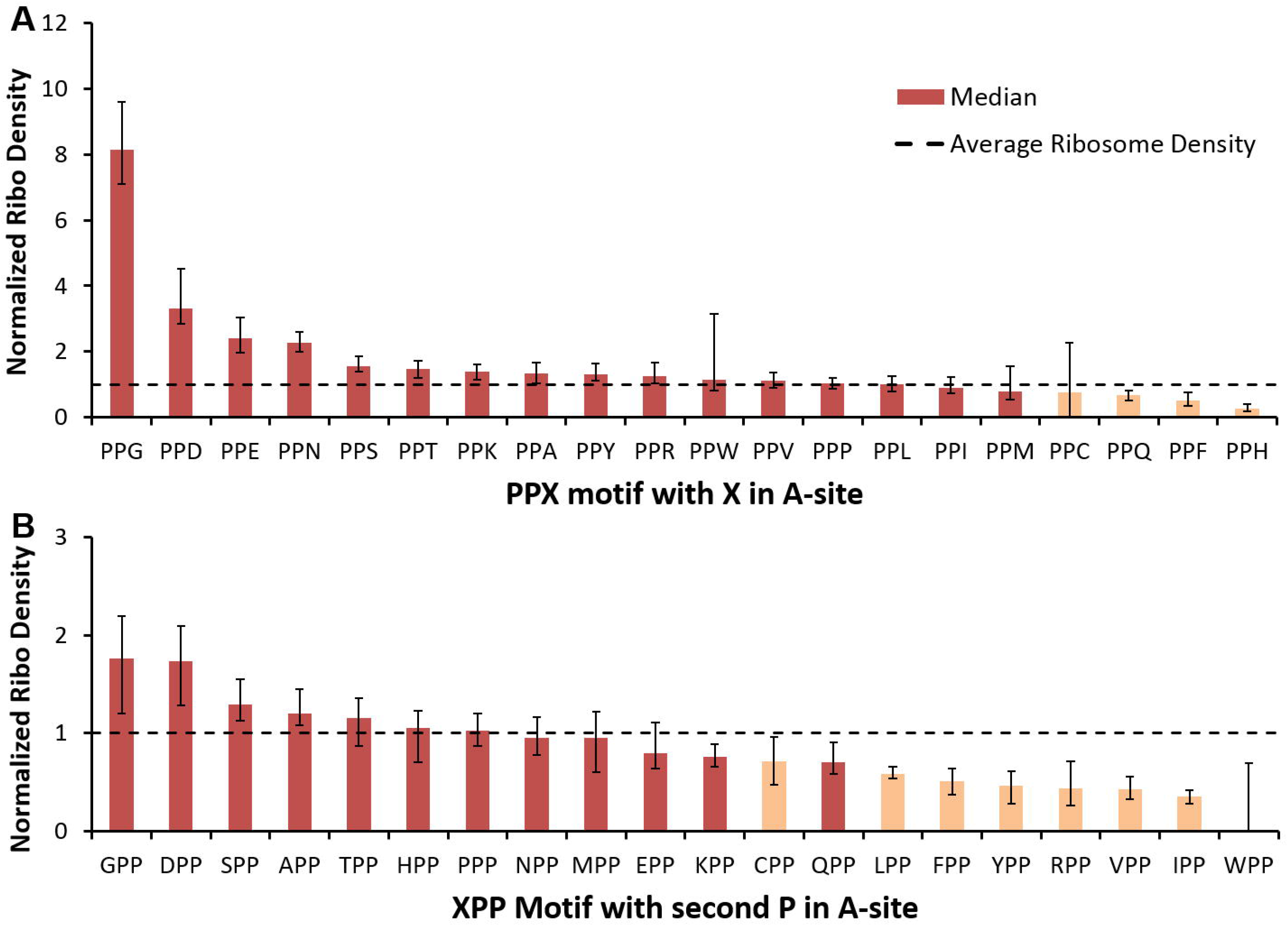
Several PPX and XPP motifs lead to ribosomal stalling in *S. cerevisiae*. The median normalized ribosome density is obtained for all instances of **(A)** PPX and **(B)** XPP motifs in which X corresponds to any one of the 20 naturally occurring amino acids. Using a permutation test, we determine if the median ribosome density is statistically significant or occurs by random chance. Statistically significant motifs are highlighted in dark red. This analysis was carried out on the Pop dataset for transcripts in which at least 50% of codon positions have reads mapped to them. Error bars are 95% Confidence Intervals for the median obtained using Bootstrapping.

A study of Ribo-Seq data of mammalian cells^49^ observed a sequence-independent translation pause when the 5^th^ codon of the transcript is in the P-site. This post-initiation pausing was also observed in an *in vitro* study of poly-phenylalanine synthesis where stalling was observed when the 4^th^ codon was in the P-site^50^. With the A-site profiles obtained using our offset tables for *S. cerevisiae* and mESCs; we also observe these pausing events when both the 4^th^ and 5^th^ codons are at the P-site (Supplementary Fig. S6).

### Greater A-site location accuracy than other methods

There is no independent experimental method to verify the accuracy of identified A-site locations using our method or any other method^4,5,52–55,6,8–10,12,41,42,51^. We argue that the well-established ribosome pausing at particular PPX sequence motifs is the best available means to differentiate the accuracy of existing methods. The reason for this is that these stalling motifs have been identified in *E.coii*^56,57^ and *S. cerevisiae*^58^ through orthogonal experimental methods (including enzymology studies and toe printing), and the exact location of the A-site during such a slowdown is known to be at the codon encoding the third residue of the motif^56^. Thus, the most accurate A-site identification method will be the one that most frequently assigns greater ribosome density to X at each occurrence of the PPX motif.

We applied this test to the strongest stalling PPX motifs, *i.e.*, PPG in *S. cerevisiae* and PPE in mESCs. In *S. cerevisiae*, the Integer Programming method yields the greatest ribosome density at the glycine codon of PPG motif when applied to both the Pooled (Fig. 6A) and Pop datasets (Supplementary Fig. S7A). Examining each occurrence of PPG in the transcriptome, we find that in a majority of instances our method assigns more ribosome density to glycine than every other method when applied to both the Pooled (Fig. 6B, Wilcoxon signed-rank test (*n* = 224), *P* < 0.0005 for all methods except Hussmann (*P* = 0.164)) and Pop datasets (Supplementary Fig. S7B, Wilcoxon signed-rank test (*n* = 35), *P* < for all methods except Hussmann (*P* = 0.026) and Ribodeblur (*P* = 0.01). The same analyses applied to mESCs at PPE motifs shows that our method outperforms the other nine methods (Figs. 6C-D) with our method assigning greater ribosome density at glutamic acid for at least 85% of the PPE motifs in our dataset as compared to all other methods (Fig. 6D, Wilcoxon signed-rank test (*n* = 104), *P* < 10^−15^ for all methods). Thus, for *S. cerevisiae* and mESCs our Integer Programming approach is more accurate than other methods in identifying the A-site on ribosome-protected fragments.

**Figure 6:**
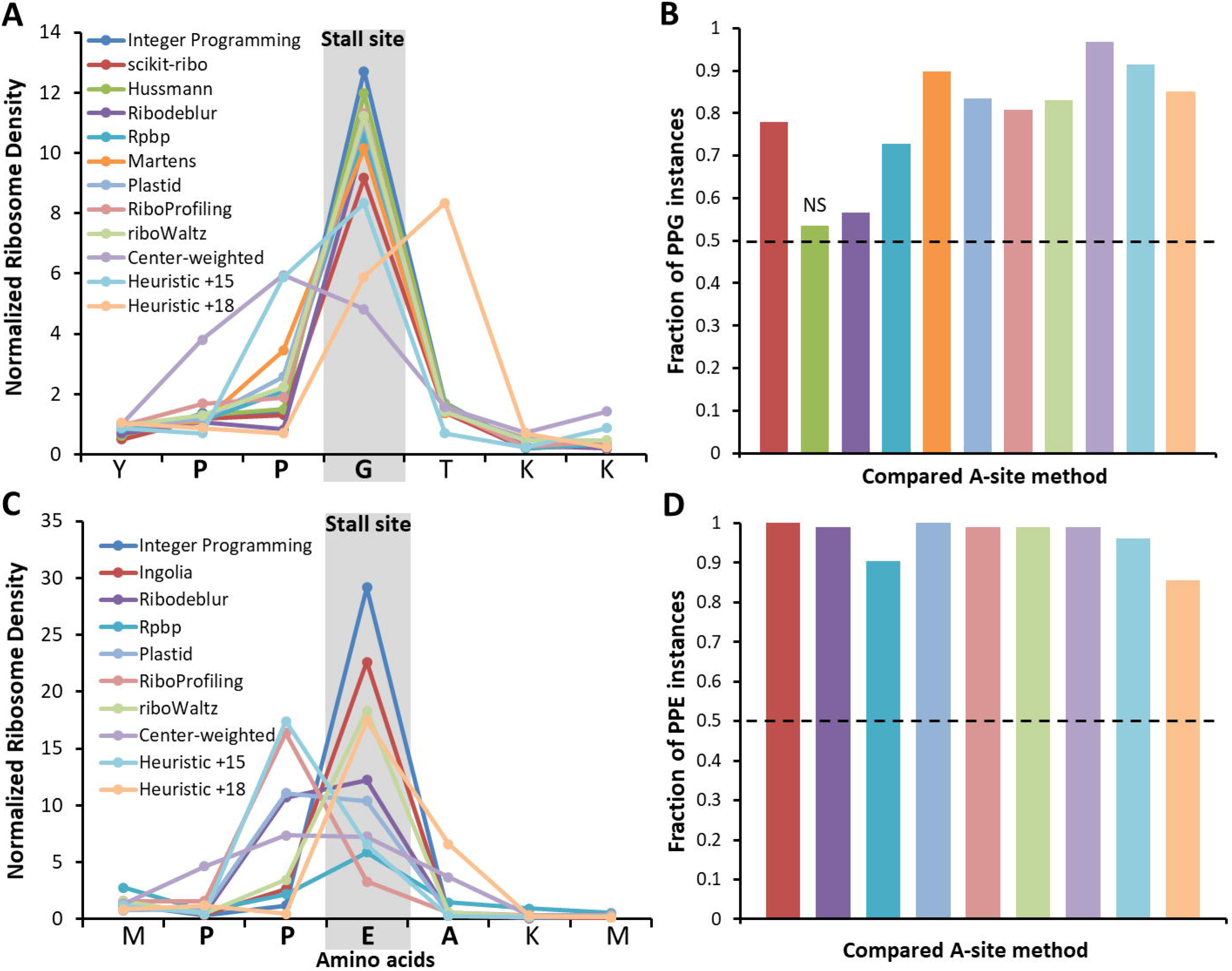
The Integer Programming algorithm correctly assigns greater ribosome density than other methods to the Glycine in PPG motifs in *S. cerevisiae* and to Glutamic acid in PPE motifs in mESCs. **(A)** Normalized ribosome density obtained using the various methods used to identify the A-site is shown for an instance of PPG motif in gene YLR374W with G at codon position 303 in the Pooled dataset of *S. cerevisiae* (see Legend and Main Text for details about methods). **(B)** The fraction of PPG instances (*n* = 224) at which the Integer Programming method yields greater ribosome density at glycine compared to every other method. The color-coding is the same as shown in the legend in panel **(A)**. Our method does better if it assigns greater ribosome density in more than half the instances (horizontal line in panel B). The Integer Programming method does better than all other methods (*P* < 0.0005) except for Hussmann, which is not statistically different (*P* = 0.164). **(C)** Normalized ribosome density is shown for an instance of PPE motif in gene uc007zma.1 with E at codon position 127 in the Pooled dataset of mouse ESCs (see Legend and main text for details about methods). **(D)** The fraction of PPE instances at which the Integer Programming method yields greater ribosome density at glutamatic acid compared to every other method. The color-coding is same as shown in the legend of panel **(C)**. The Integer Programming method does better than all other methods (*P* > 10^−15^) in accurately assigning ribosome density to Glutamic Acid in PPE motifs (*n* = 104). For both analyses, two-sided *p*-values were calculated using the Wilcoxon signed rank test. Error bars represent the 95% Confidence Interval about the median calculated using Bootstrapping.

A large number of molecular factors influence codon translation rates and ribosome density along transcripts^59^. One factor is the cognate tRNA concentration, as codons decoded by cognate tRNA with higher concentrations should have on average lower ribosome densities^15,16,60^. Therefore, as an additional qualitative test, we expect that the most accurate A-site method will yield the largest anti-correlation between the ribosome density at a codon and its cognate tRNA concentration. This test is only qualitative as the correlation between codon ribosome-density and cognate tRNA concentration may be affected by other factors, including codon usage and reuse of recharged tRNAs in the vicinity of the ribosome influence the relationship^61,62^. Using tRNA abundances previously estimated from RNA-Seq experiments on *S. cerevisiae*^16^, we find that our Integer Programming method yields the largest anti-correlation compared to the eleven other methods considered (Supplementary Table S10), further supporting the accuracy of our method. We were unable to run this test in mESCs as measurements of tRNA concentration have not been reported in the literature.

## Discussion

We have introduced a method to determine the A- and P-site locations on ribosome-protected mRNA fragments, and shown that it is more accurate than other methods in correctly assigning ribosome density to the glycine residue in PPG motifs and glutamic acid residue in PPE motifs, which are strong translation-stalling sites in *S. cerevisiae* and mESCs, respectively. Our method is unique amongst existing methods because it (*i*) uses a probabilistic approach to identify the A-site location through Integer Programming optimization and (*ii*) has an objective function rooted in the biology of translation - meaning that its optimization enforces the fact that the A-site location of most reads must have been between the second and stop codons of the CDSs. To be sure, several methods use biological features to assign the A-site (such as having more reads around the start and stop codons than in the UTR^2,11^). However, ours is the only method that also utilizes feature (*i)*, which is beneficial because the stochastic nature of mRNA cleavage during the digestion-step of Ribo-Seq necessitates a probabilistic perspective. Our method is not entirely probabilistic since we have to set thresholds and apply a secondary criterion to arrive at a final offset value. These measures are unavoidable due to the variability in coverage between different genes. However, we find that the results are robust to variation in thresholds and mostly consistent across different Ribo-Seq datasets. Hence, the respective A-site offset tables provided for *S. cerevisiae* (Table 1) and mouse embryonic stem cells (Supplementary Table S8) can be applied to any dataset from these organisms.

Noteworthy about our test for accuracy is that it is based on results from orthogonal experimental techniques. The stalling of translation at glycine in PPG motifs is well-documented^33,47,56–58^ and in *S. cerevisiae* the Integer Programming method assigns higher Ribo-Seq reads at the glycine codon at most instances of PPG compared to other A-site methods. In mESCs PPE is the strongest stalling motif^11^. The Integer Programming method outperforms other methods by assigning, on average, 1.76 times more reads at the glutamic acid codon compared to other methods. These results indicate that the Integer Programming method presented in this study is more accurate than existing methods. One reason for this increase in accuracy, among many possible reasons, may be that most methods only use reads from around the start codon, while our method uses reads from around both the start and stop codons.

A potential point of confusion may arise from the distributions shown in Fig. 3 in which there are two highly probable offset values, raising the question of whether or not there are multiple A-site locations for a given fragment size and frame. In almost all fragment length and frame combinations, there is one unique most probable A-site location, but this ambiguity can arise from poor read coverage on a gene or stochastic fluctuations in the extent of digestion on one side of an mRNA fragment compared to the other. Consider fragment size 28 in frame 1. In the Pop data set (top, middle panel of Fig. 3A), approximately half of the genes have Δ= 15 nt, while the others have Δ= 18 nt, meaning the A-site could be at either location. When we increase the read coverage of the genes, however, we see that the vast majority of the offsets shift to 15 nt (bottom, middle panel in Fig. 3B). Thus, the original A-site ambiguity was not due to multiple, equally possible A-site locations, but rather the true A-site location was hard to detect without better coverage. Consider another example. For *S* = 27 and *F* = 1 we observe in Fig. 3A that 8% of genes have an optimal Δ= 0, seemingly suggesting that the A-site is located at the 5□-end on a subset of fragments. Spot-checking the ribosome profiles of these genes, we find that these genes contain no reads in the 27 nt region upstream of the second codon and 27 nt upstream of the stop codon (data not shown). Thus, the values of *T*(Δ|*i,S,F*) for all Δwere equal and the optimal Δwas arbitrarily assigned a value of 0. In the higher coverage Pooled dataset, however, there are only 2% of genes with optimal Δ= 0 for S’ = 27 and *F* = 1. Hence, as we increase coverage, the proportion of genes with spurious offsets decreases. Thus, offsets away from the most probable offset arise from sampling issues, not from multiple A-site locations. This result is also seen in the analysis of the artificial Ribo-Seq data where our algorithm correctly predicts the true offsets for a majority of (*S, F*) combinations while ambiguous offsets occur only for those (*S,F*) combinations with the lowest read coverage.

We note that we set a threshold of 70% to determine a most-probable offset for each fragment size and reading frame and demonstrated that the results are robust to variation with this threshold (Supplementary Table S5). Therefore, the A-site assignments reported in Table 1 represent the most likely location of the A-site relative to the 5□ end of mRNA fragments produced from Ribo-Seq experiments on *S. cerevisiae*.

Some (*S,F*) combinations (such as *S* = 32 and *F* = 0, in Table 1) appear to be inherently ambiguous, that is, increasing their coverage does not lead to a unique A-site assignment (Fig. 4D). We do not know the reason for this result, but we speculate that these are situations where there are truly multiple equally probable A-site locations. Another possibility is that the ribosome adopts different conformations in these situations that result in different read lengths and offsets, leading to ambiguity^14^. The important point is that the A-site cannot be accurately assigned in these situations. We therefore recommend that researchers discard reads from these (*S, F*) combinations to minimize chances of erroneous A-site assignments. We believe it will have negligible effect on the A-site profiles since these combinations contribute only 2.9% of total reads in the Pooled dataset.

We have found that the Integer Programming algorithm is sensitive to reads arising from outside the boundaries of annotated CDS regions from non-canonical sources like upstream ORFs (uORFs) or Internal Ribosome Entry Sites (IRES). Specifically, applying our method to Ribo-Seq data from mESCs yielded few unique offsets. It was only after removing genes that had multiple translation initiation sites, some arising from uORFs, that the number of unique offsets increased more than four-fold. The reason for this improvement was that by removing the uORFs, our method’s assumption was met that the reads within 40 nt of the start codon only arise from the annotated CDS. Our method was not able to identify any unique offsets in *E. coli* Ribo-Seq data even after we controlled for multiple translation initiation sites. We observed in *E. coli* a high enrichment of reads before the start codon after applying the conventional 12 nt offset from 3 end^33^ (Supplementary Fig. S4) which we speculate may be due to protection of mRNA segments involved in binding of the Shine-Dalgarno sequence to the ribosome^63^ and could limit the accuracy of our method.

The next best method to the Integer Programming method is the Hussmann approach^12^. Besides more frequently assigning greater ribosome density to glycine in PPG motifs and exhibiting strong correlation with cognate tRNA abundances, the Integer Programming method is also superior because it provides greater statistical power and is based on biological features of translation rather than heuristic assumptions. Specifically, Hussmann’s method only uses reads that are 28, 29 and 30 nt in length, whereas our method uses reads between 24 to 34 nt in length. This greater coverage results in greater statistical power for our method. Hussmann’s method uses a nearest-neighbour heuristic to determine frame-specific offsets of +14, 15 or 16 for lengths 28 and 29 and offset of +15, 16 or 17 for length 30, whereas our method is based on the feature that the A-site be located within the CDS. The reason Hussman’s method yields comparable results is that its offset table is highly similar to Table 1. If the reading frame is maintained after applying the offset from the 5□ end, then 8 out of 9 of Hussmann’s offsets are the same as in Table 1 with the 9^th^ offset of (29,1) being ambiguous in our method.

Our method preserves the original 3 nt periodicity found in the original 5□-end aligned mRNA fragments. Therefore, it is not designed for detecting frame-shifting, translation of upstream ORFs, or novel short peptides. Nevertheless, correct assignment of reads to the A-site codon is essential in a variety of other analyses, such as determining translation kinetics, and our method provides the most accurate assignment of ribosome density compared to other methods (Fig. 6 and Supplementary Table S10).

In summary, we have created a method for A-site identification that is more accurate than existing methods in *S. cerevisiae* and mouse embryonic stem cells, utilizes a fundamental feature of translation to identify the A-site, and have revealed how the A-site location changes based on the size of the mRNA fragment and its frame. By increasing the accuracy and range of fragment sizes for which the A-site can be identified, our approach can help future studies to measure translation elongation properties at the length scale of individual codons.

## Acknowledgements

We thank the members of the O’Brien Lab for critical feedback on the manuscript. PS is supported by a Borysiewicz Biomedical Fellowship from the University of Cambridge. This work was supported by the research grant from the National Science Foundation ABI grant 1759860 to EPO.

## Author Contributions

PS, PC and EPO conceived the study. NA, PS and EPO designed the computational analyses. PC, MV, CMD contributed to design of the computational analyses. NA and PS analyzed the data. NA and EPO wrote the manuscript. All authors reviewed and commented on the manuscript.

## Data Availability

All source code is made available on the GitHub repository https://github.com/nabeel1990/Asite_IP_method

