## Supplementary Information for "Identifying A- and P-site locations on ribosome-protected mRNA fragments using Integer Programming"

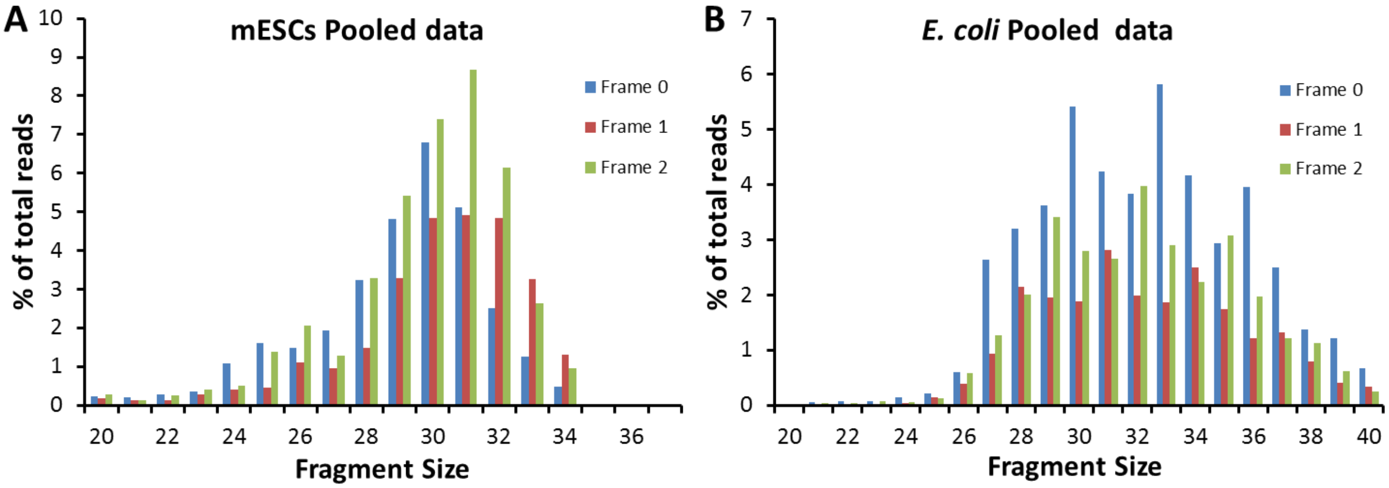


**Supplementary Figure S1:** Fragment size distribution in **(A)** Pooled Ribo-Seq data in mouse embryonic stem cells (mESCs) and **(B)** Pooled Ribo-Seq data in *Escherichia coli.*

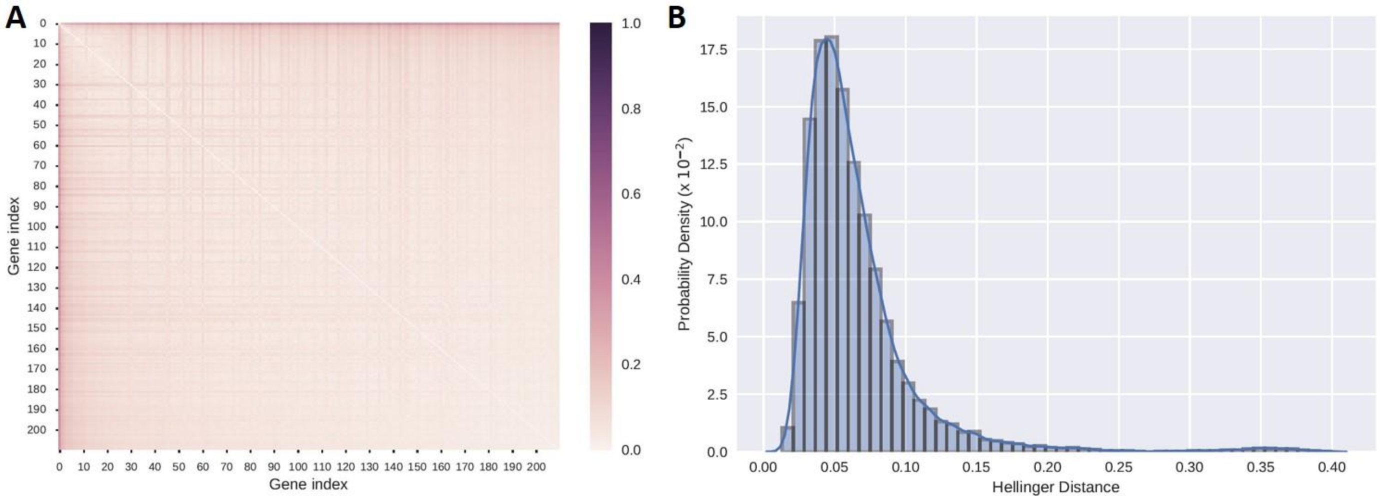


**Supplementary Figure S2: Pairwise comparison of fragment-size and frame distributions between genes in *S. cerevisiae*. (A)** The heat map reports the pairwise Hellinger distance[1] between the probability densities of the fragment-size and frame distributions of individual genes. Only genes in the Pooled data set that have at least 1 read per codon for fragment sizes between 24 and 34 nt were analyzed, resulting in 210 genes in this analysis. **(B)** The probability density distribution of Hellinger distances reported in (A). The Hellinger distance metric is bound between (0, 1); 0 indicates identical distributions; while 1 indicates the distributions are divergent. All pairwise Hellinger distances are less than 0.45 and only 11% of pairwise distances are greater than 0.1. Hence, the distribution of reads of different fragment sizes and frames are highly similar and therefore exhibit very little dependence on gene identity.

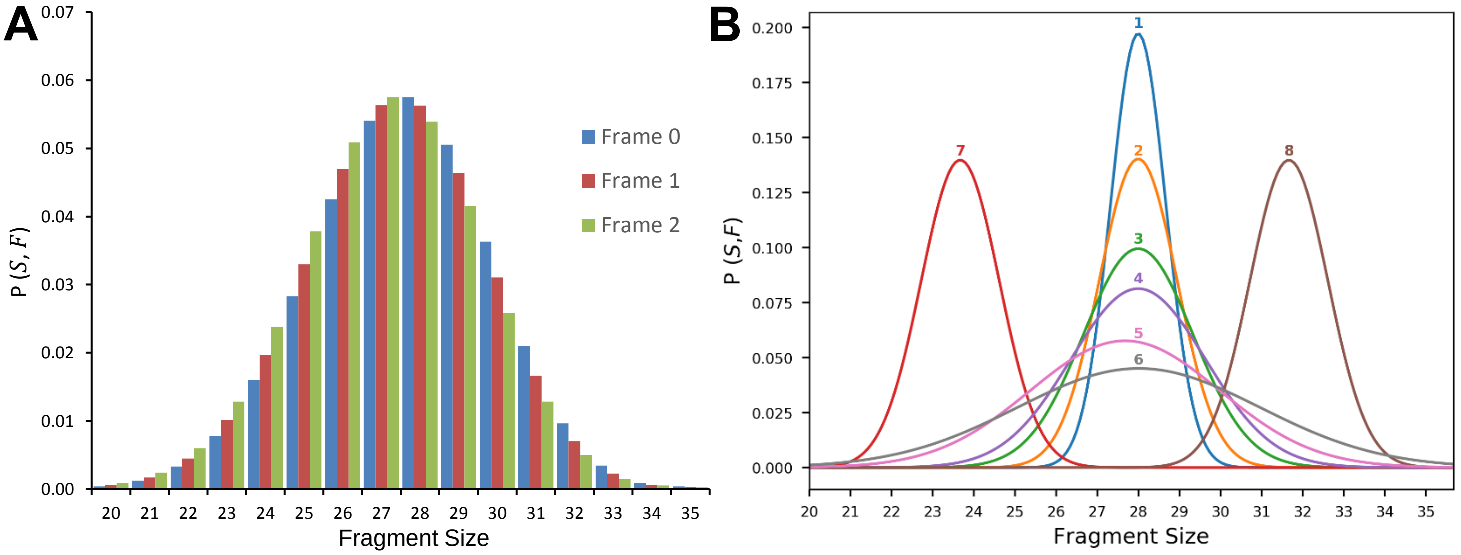


**C**

| Read length Distribution | Input offset table | | | |
|---|---|---|---|---|
| | Constant offset of 15 | Constant offset of 18 | Mixed offsets of 12 and 18 | Top offsets from experimental data |
| 1 | 100% | 93% | 100% | 93% |
| 2 | 95% | 95% | 100% | 95% |
| 3 | 96.5% | 100% | 96.5% | 100% |
| 4 | 100% | 100% | 100% | 97% |
| 5 | 100% | 96% | 100% | 98% |
| 6 | 100% | 100% | 100% | 100% |
| 7 | 100% | 100% | 100% | 100% |
| 8 | 95% | 95% | 95% | 100% |

**Supplementary Figure S3: Integer Programming algorithm correctly reproduces the true A-site offsets from Artificial Ribo-Seq data. (A)** An example of a Poisson distribution with mode at $(S, F) = (28, 0)$ and a variance $\lambda = 48$ that was used to generate artificial ribosome-protected fragments (see Methods for details). The reads generated from this distribution are subjected to the Integer Programming algorithm for four different input offset tables. These input offset tables are shown in Table S6. **(B)** Six read length distributions with their mode at $(28, 0)$ were generated with Poisson variances $\lambda = 4, 8, 16, 24, 48, 80$ and labeled 1 through 6, respectively. The distribution in (A) is the distribution labeled 5 in (B). Two more distributions were generated with variance $\lambda = 8$ but with modes at $(24, 0)$ and $(32, 0)$ and labelled as 7 and 8, respectively. **(C)** The percentage of offsets that the Integer Programming correctly identifies in the artificial Ribo-Seq data created based on the eight read length distributions shown in (B) and for each of the four different input offset tables (see Methods for details) used to generate the artificial Ribo-Seq reads. The four input offset tables and the corresponding output offset tables generated by the Integer Programming algorithm for distribution 5 is shown in Table S6.

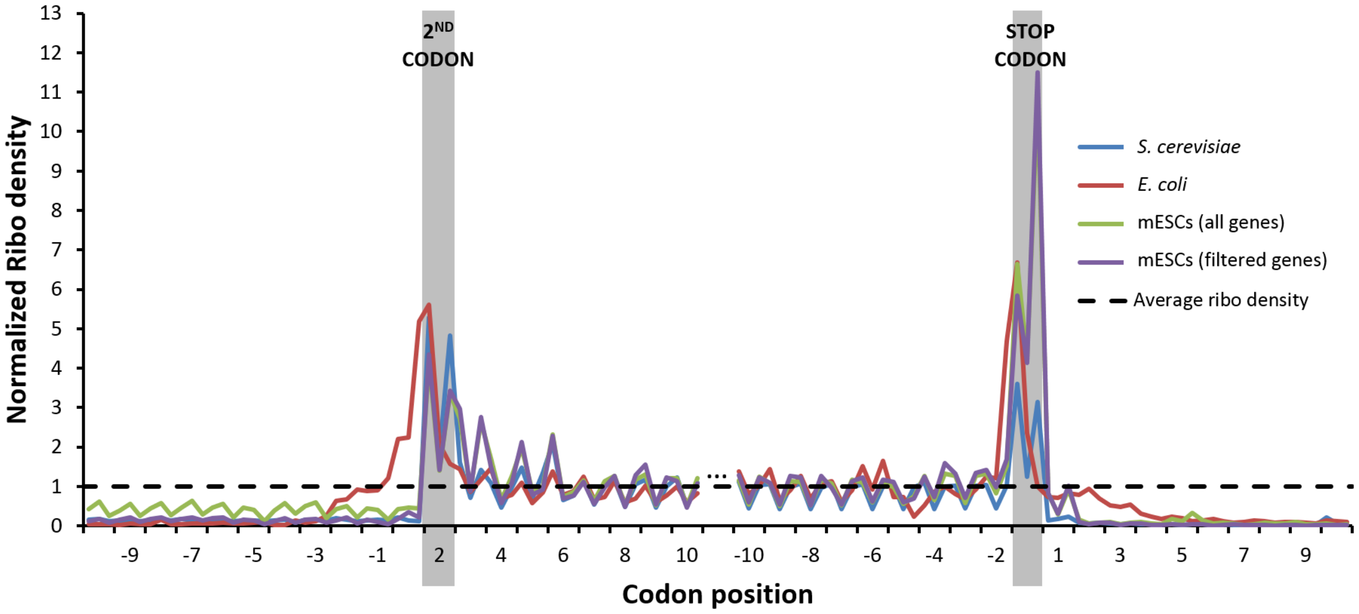


**Supplementary Figure S4: Meta-gene analysis in Pooled Ribo-Seq data reveal excess ribosome density in *E.coli* genes beyond CDS regions.** A-site profiles are obtained for *S. cerevisiae* using unique offsets from Table 1 obtained after application of Integer Programming algorithm. For *E. coli*, we use a constant offset of 12 nt from 3′ end as used by Woolstenhulme and co-workers [2]. For mESCs, we use the unique offsets from Table S6. We plot the meta-gene profiles for all genes dataset as well as for the subset of filtered genes containing only single isoforms with one translation start site.

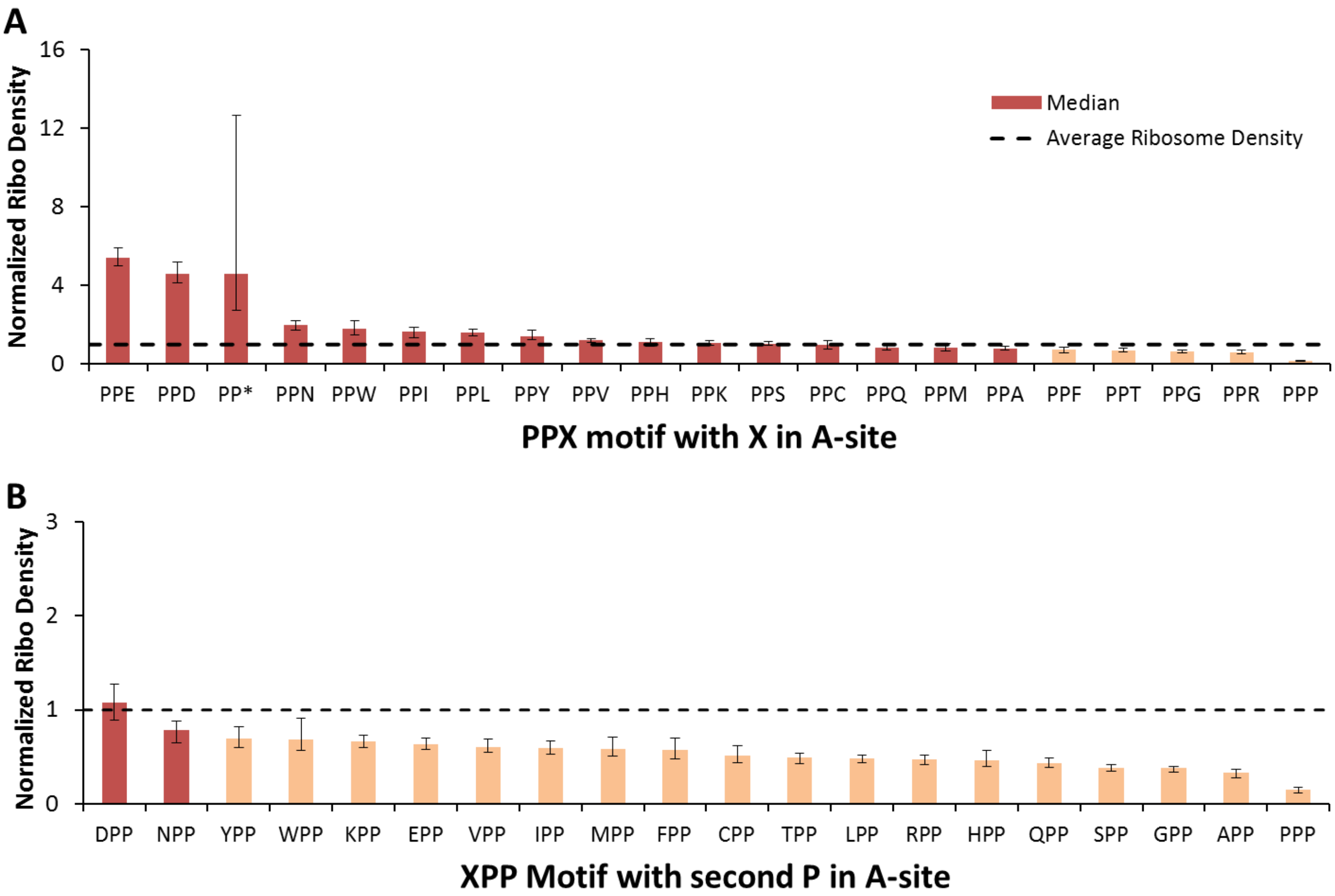


**Supplementary Figure S5: Stalling at PPE and PPD motifs are reproduced in mESCs.** The median normalized ribosome density is obtained for all instances of **(A)** PPX and **(B)** XPP motifs in which X corresponds to any one of the 20 naturally occurring amino acids (and stop codon for instance PP*). Using a permutation test, we determine if the median ribosome density is statistically different from the average ribosome density. Statistically significant motifs are highlighted in dark red. This analysis was carried out on the Pooled dataset for transcripts in which at least 50% of codon positions have reads mapped to them. Error bars are 95% Confidence Intervals for the median obtained using Bootstrapping [3].

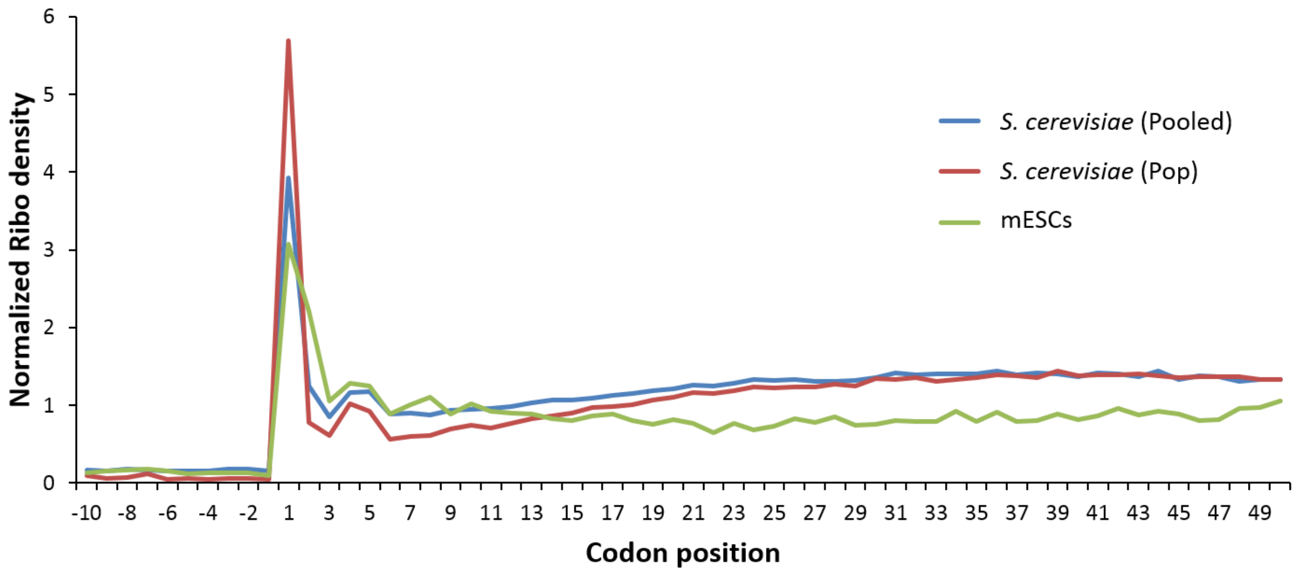


**Supplementary Figure S6: Sequence-independent translational pause observed post-initiation in *S. cerevisiae* and mESCs.** Meta-gene analysis at the codon level with reads mapped to the P-site are shown. There is a mild but distinct pausing of translation when the 4$^{th}$ and 5$^{th}$ codons are in the P-site. This effect is seen in both Pooled and Pop datasets of *S. cerevisiae* as well as the filtered genes dataset of mouse embryonic stem cells (mESCs).

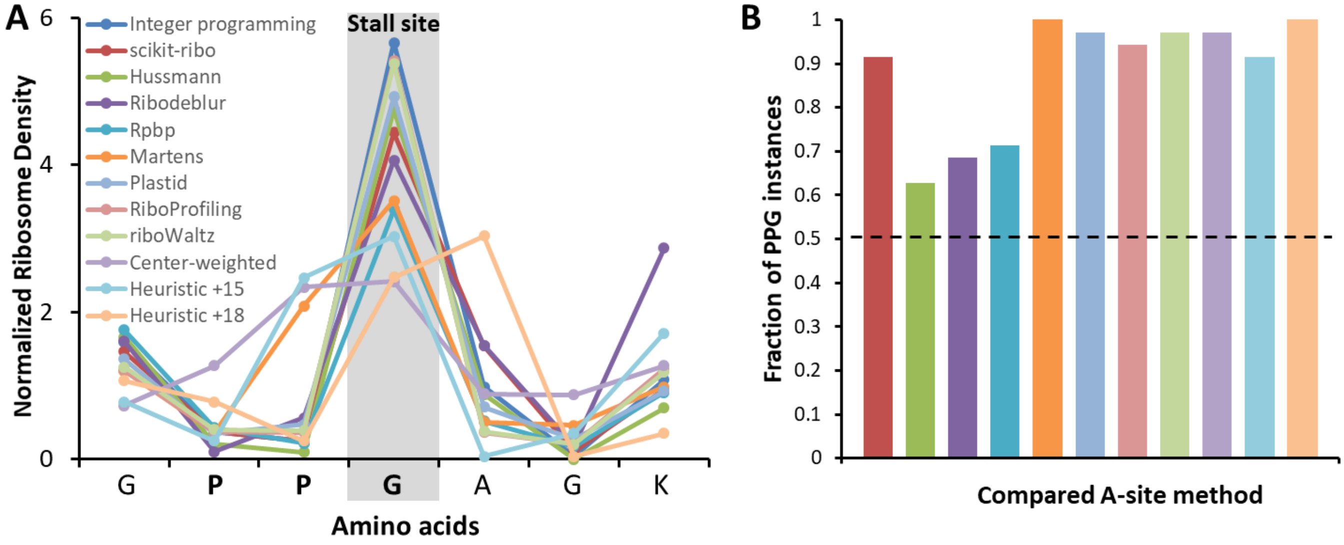


**Supplementary Figure S7: The Integer Programming algorithm correctly assigns greater ribosome density to the Glycine residue in PPG motifs than other methods in *S. cerevisiae*.** **(A)** Normalized ribosome density at the predicted A-site using different methods to determine A-site is shown for an instance of the PPG motif in gene YDR226W with the Glycine of the PPG motif is at codon position 16, for Pop dataset in *S. cerevisiae*. **(B)** The fraction of PPG instances ($n = 35$) at which Integer Programming method yields greater ribosome density at glycine against the compared method. The color-coding is same as shown in the legend of panel (A). Our method does better if it assigns greater ribosome density in more than half the instances (horizontal line in panel B). Integer Programming yields significantly higher ribosome density at G in the PPG motifs than all other methods (For Hussmann $P = 0.026$, for ribodeblur $P = 0.01$ and for others $P < 10^{-5}$). Two-sided *P*-values were calculated using the Wilcoxon signed rank test. Error bars are 95% Confidence Interval about the median calculated using Bootstrapping [3].

**Supplementary Table S1:** Publicly available datasets used in the study.

| Dataset (first author name) | Year of publication | Number of replicates* | GEO Study | Accession numbers of samples used |
|---|---|---|---|---|
| *Saccharomyces cerevisiae* | | | | |
| Pop | 2014 | 1 | GSE63789 | GSM1557447 |
| Guydosh | 2014 | 1 | GSE52968 | GSM1279568 |
| Jan | 2014 | 1 | GSE61012 | GSM1495525 |
| Williams | 2014 | 1 | GSE61011 | GSM1495503 |
| Gerashchenko | 2014 | 1 | GSE59573 | GSM1439584 |
| Gardin | 2014 | 2 | GSE51164 | GSM1239959, GSM1239960 |
| Lareau | 2014 | 3 | GSE58321 | GSM1406453, GSM1406454, GSM1406455 |
| Nedialkova | 2015 | 3 | GSE67387 | GSM1646015, GSM1646016, GSM1646017 |
| Young | 2015 | 1 | GSE69414 | GSM1700885 |
| Weinberg | 2016 | 1 | GSE53268 | GSM1289257 |
| Nissley | 2016 | 2 | GSE75322 | GSM1949550, GSM1949551 |
| Mouse embryonic stem cells (mESCs) | | | | |
| Ingolia | 2011 | 1 | GSE30839 | GSM765298 |
| Hurt | 2013 | 1 | GSE41785 | GSM1024298 |
| *Escherichia coli* | | | | |
| Li | 2012 | 2 | GSE35641 | GSM872393, GSM872394 |
| Li | 2014 | 1 | GSE53767 | GSM1300279 |
| Woolstenhulme | 2015 | 2 | GSE64488 | GSM1572273, GSM1572275 |

* For datasets with more than one replicate, all replicates were used to create the Pooled dataset.

**Supplementary Table S2:** Number of genes for the various fragment size and frame combinations that meet the criteria of at least 1 read per codon on average in the Pop and Pooled datasets of *S. cerevisiae*.

| Fragment size | Pop dataset | | | | Pooled dataset | | |
|---|---|---|---|---|---|---|---|
| | Frame 0 | Frame 1 | Frame 2 | | Frame 0 | Frame 1 | Frame 2 |
| **20** | 34 | 7 | 57 | | 156 | 50 | 98 |
| **21** | 45 | 60 | 41 | | 224 | 129 | 139 |
| **22** | 99 | 55 | 48 | | 234 | 115 | 199 |
| **23** | 73 | 47 | 95 | | 162 | 107 | 290 |
| **24** | 58 | 91 | 72 | | 161 | 352 | 193 |
| **25** | 155 | 69 | 55 | | 647 | 251 | 194 |
| **26** | 105 | 64 | 175 | | 481 | 241 | 916 |
| **27** | 159 | 255 | 161 | | 1096 | 2213 | 878 |
| **28** | 1081 | 248 | 333 | | 4487 | 1861 | 1468 |
| **29** | 850 | 330 | 1437 | | 3919 | 1504 | 4474 |
| **30** | 528 | 876 | 1139 | | 3184 | 3041 | 3835 |
| **31** | 276 | 610 | 643 | | 2089 | 2251 | 3164 |
| **32** | 70 | 279 | 181 | | 799 | 1897 | 1789 |
| **33** | 39 | 82 | 9 | | 474 | 1076 | 322 |
| **34** | 1 | 2 | 0 | | 237 | 194 | 71 |
| **35** | 0 | 0 | 0 | | 58 | 40 | 33 |

**Supplementary Table S3:** Number of genes in the combination of fragment size and frame meeting the criteria of at least 1 read per codon on average in mESCs and *E. coli* Pooled datasets. The mESCs Pooled dataset consist of genes that are single isoform and have only one defined translation initiation site.

| Fragment size | mESCs Pooled dataset | | | | *E. coli* Pooled dataset | | |
|---|---|---|---|---|---|---|---|
| | Frame 0 | Frame 1 | Frame 2 | | Frame 0 | Frame 1 | Frame 2 |
| **20** | 8 | 7 | 10 | | 313 | 243 | 330 |
| **21** | 10 | 0 | 1 | | 440 | 270 | 377 |
| **22** | 10 | 1 | 10 | | 532 | 416 | 431 |
| **23** | 15 | 15 | 18 | | 610 | 471 | 645 |
| **24** | 41 | 20 | 19 | | 806 | 471 | 655 |
| **25** | 61 | 19 | 52 | | 816 | 610 | 603 |
| **26** | 52 | 43 | 75 | | 765 | 625 | 742 |
| **27** | 73 | 38 | 45 | | 952 | 681 | 825 |
| **28** | 126 | 55 | 119 | | 1001 | 849 | 868 |
| **29** | 187 | 125 | 208 | | 988 | 840 | 981 |
| **30** | 230 | 191 | 257 | | 1072 | 791 | 956 |
| **31** | 197 | 192 | 280 | | 1042 | 898 | 916 |
| **32** | 103 | 187 | 237 | | 994 | 823 | 993 |
| **33** | 47 | 125 | 108 | | 1060 | 761 | 943 |
| **34** | 17 | 55 | 45 | | 1008 | 842 | 856 |
| **35** | 0 | 0 | 0 | | 891 | 740 | 924 |
| **36** | 0 | 0 | 0 | | 943 | 598 | 799 |
| **37** | 0 | 0 | 0 | | 827 | 618 | 625 |
| **38** | 0 | 0 | 0 | | 640 | 494 | 596 |
| **39** | 0 | 0 | 0 | | 588 | 323 | 461 |
| **40** | 0 | 0 | 0 | | 440 | 288 | 278 |

**Supplementary Table S4:** Initial offset tables after application of Integer Programming algorithm to Pop and Pooled datasets in *S. cerevisiae.*

| Fragment size | Pop dataset | | | Pooled dataset | | |
|---|---|---|---|---|---|---|
| | Frame 0 | Frame 1 | Frame 2 | Frame 0 | Frame 1 | Frame 2 |
| **20** | 0/6 | ND* | 9/0 | 6/15 | 6/0 | 9/0 |
| **21** | 15/6 | 9/15 | 9/0 | 15/6 | 9/0 | 9/18 |
| **22** | 9/0 | 9/18 | 18/9 | 9/15 | 9/0 | 18/9 |
| **23** | 9/15 | 12/15 | 18/12 | 9/15 | 9/18 | 18/12 |
| **24** | 15/12 | 12/18 | 18/12 | 15/9 | 12/15 | 18/12 |
| **25** | 12/15 | 12/18 | 18/12 | 15/12 | 12/15 | 18/12 |
| **26** | 15 /12 | 15/18 | 18/15 | 15 /12 | 12/15 | 18/15 |
| **27** | **15** | 15/18 | 18/15 | **15** | **15** | 18/15 |
| **28** | **15** | 18/15 | **18** | **15** | **15** | 18/15 |
| **29** | **15** | 18/15 | **18** | **15** | 15/18 | **18** |
| **30** | **15** | **18** | **18** | **15** | 18/15 | **18** |
| **31** | 18/15 | **18** | **18** | **15** | **18** | **18** |
| **32** | 18/15 | **18** | **18** | 18/15 | **18** | **18** |
| **33** | 18/0 | 18/0 | ND | 18/15 | **18** | 18/15 |
| **34** | ND | ND | ND | 18/15 | 18/15 | 18/21 |
| **35** | ND | ND | ND | 18/15 | 15/18 | 21/18 |

**Supplementary Table S5:** For unique offsets described in Table 1, the robustness to variation in parameters and consistency across different Ribo-Seq datasets are described with additional sub columns. The two sub columns in the top row refers to the unique offsets being sensitive (S) or robust (R) to parameter variation. Namely, the change in threshold from 60% to 80% to classify the most probable offset as unique (left sub column) and variation in threshold of the secondary selection criterion $R(1) < a * Mean\big(R(2), R(3), R(4)\big)$ where $a$ ranges from 1 to $\frac{1}{10}$ (right sub column). The bottom row specifies the consistency of the unique offset across individual Ribo-Seq datasets. For example, for fragment size 27 in frame 0, 15 is the unique offset which is sensitive (S in left sub column) to a change in threshold from 60% to 80% and robust (R in right sub column) to change in secondary selection parameter $a$ from 1 to $\frac{1}{10}$. It is also consistent in 9 out of 12 datasets for which we have more than 10 genes meeting our filtering criteria.

| **Fragment Size** | **Frame 0** | | | | **Frame 1** | | | | **Frame 2** | | | |
|---|---|---|---|---|---|---|---|---|---|---|---|---|
| **24** | **15** | R | R | 4 of 4 | 15/12 | | | | 18/12 | | | |
| **25** | **15** | S | R | 5 of 7 | 12/15 | | | | **18** | S | S | 4 of 4 |
| **26** | 15/12 | | | | 18/15 | | | | 18/15 | | | |
| **27** | **15** | S | R | 9 of 12 | **15** | S | R | 7 of 12 | **18** | R | R | 6 of 9 |
| **28** | **15** | R | R | 14 of 17 | **15** | S | R | 10 of 13 | **18** | R | R | 10 of 12 |
| **29** | **15** | R | R | 14 of 15 | 15/18 | | | | **18** | R | R | 15 of 16 |
| **30** | **15** | R | R | 15 of 15 | **18** | R | R | 12 of 16 | **18** | R | R | 16 of 16 |
| **31** | **15** | S | R | 12 of 13 | **18** | R | R | 13 of 15 | **18** | R | R | 15 of 16 |
| **32** | 18/15 | | | | **18** | R | R | 11 of 11 | **18** | R | R | 9 of 10 |
| **33** | **18** | S | S | 6 of 6 | **18** | R | R | 7 of 7 | **18** | S | S | 4 of 4 |
| **34** | **18** | R | R | 2 of 2 | **18** | R | S | 2 of 2 | 18/21 | | | |

**Supplementary Table S6:** Input A-site offset tables used in the creation of artificial Ribo-Seq data (Top row, see Methods). Offset A-site tables (Bottom row) output by the Integer Programming method when applied to artificial Ribo-Seq data constructed using the input tables (Top) and P$(S,F)$ distribution with mode $(28,0)$ and variance $\lambda = 48$ (Distribution 5 in Fig S3).

**Input Offset tables**

| Fragment size | Constant offset of 15 | | | Constant offset of 18 | | | Mixed offsets of 12 and 18 | | | Top offsets from exp. data | | |
|---|---|---|---|---|---|---|---|---|---|---|---|---|
| | Frame 0 | Frame 1 | Frame 2 | Frame 0 | Frame 1 | Frame 2 | Frame 0 | Frame 1 | Frame 2 | Frame 0 | Frame 1 | Frame 2 |
| 20 | 15 | 15 | 15 | 18 | 18 | 18 | 12 | 12 | 12 | 6 | 6 | 9 |
| 21 | 15 | 15 | 15 | 18 | 18 | 18 | 12 | 12 | 12 | 15 | 9 | 9 |
| 22 | 15 | 15 | 15 | 18 | 18 | 18 | 12 | 12 | 12 | 15 | 6 | 18 |
| 23 | 15 | 15 | 15 | 18 | 18 | 18 | 12 | 12 | 12 | 15 | 18 | 18 |
| 24 | 15 | 15 | 15 | 18 | 18 | 18 | 12 | 12 | 12 | 15 | 15 | 12 |
| 25 | 15 | 15 | 15 | 18 | 18 | 18 | 12 | 12 | 12 | 15 | 12 | 18 |
| 26 | 15 | 15 | 15 | 18 | 18 | 18 | 12 | 12 | 12 | 15 | 15 | 18 |
| 27 | 15 | 15 | 15 | 18 | 18 | 18 | 12 | 12 | 12 | 15 | 15 | 18 |
| 28 | 15 | 15 | 15 | 18 | 18 | 18 | 18 | 18 | 18 | 15 | 15 | 18 |
| 29 | 15 | 15 | 15 | 18 | 18 | 18 | 18 | 18 | 18 | 15 | 15 | 18 |
| 30 | 15 | 15 | 15 | 18 | 18 | 18 | 18 | 18 | 18 | 15 | 18 | 18 |
| 31 | 15 | 15 | 15 | 18 | 18 | 18 | 18 | 18 | 18 | 15 | 18 | 18 |
| 32 | 15 | 15 | 15 | 18 | 18 | 18 | 18 | 18 | 18 | 15 | 18 | 18 |
| 33 | 15 | 15 | 15 | 18 | 18 | 18 | 18 | 18 | 18 | 18 | 18 | 18 |
| 34 | 15 | 15 | 15 | 18 | 18 | 18 | 18 | 18 | 18 | 18 | 18 | 18 |
| 35 | 15 | 15 | 15 | 18 | 18 | 18 | 18 | 18 | 18 | 18 | 18 | 18 |

**Output Offset tables**

| Fragment size | Constant offset of 15 | | | Constant offset of 18 | | | Mixed offsets of 12 and 18 | | | Top offsets from exp. data | | |
|---|---|---|---|---|---|---|---|---|---|---|---|---|
| | Frame 0 | Frame 1 | Frame 2 | Frame 0 | Frame 1 | Frame 2 | Frame 0 | Frame 1 | Frame 2 | Frame 0 | Frame 1 | Frame 2 |
| 20 | 15 | 15 | 15 | 18 | 18 | 18 | 12 | 12 | 12 | 6 | 6 | 9 |
| 21 | 15 | 15 | 15 | 18 | 18 | 18 | 12 | 12 | 12 | 15 | 9 | 9 |
| 22 | 15 | 15 | 15 | 18 | 18 | 18 | 12 | 12 | 12 | 15 | 6 | 18 |
| 23 | 15 | 15 | 15 | 18 | 18 | 18 | 12 | 12 | 12 | 15 | 18 | 18 |
| 24 | 15 | 15 | 15 | 18 | 18 | 18 | 12 | 12 | 12 | 15 | 15 | 12 |
| 25 | 15 | 15 | 15 | 18 | 18 | 18 | 12 | 12 | 12 | 15 | 12 | 18 |
| 26 | 15 | 15 | 15 | 18 | 18 | 18 | 12 | 12 | 12 | 15 | 15 | 18 |
| 27 | 15 | 15 | 15 | 18 | 18 | 18 | 12 | 12 | 12 | 15 | 15 | 18 |
| 28 | 15 | 15 | 15 | 18 | 18 | 18 | 18 | 18 | 18 | 15 | 15 | 18 |
| 29 | 15 | 15 | 15 | 18 | 18 | 18 | 18 | 18 | 18 | 15 | 15 | 18 |
| 30 | 15 | 15 | 15 | 18 | 18 | 18 | 18 | 18 | 18 | 15 | 18 | 18 |
| 31 | 15 | 15 | 15 | 18 | 18 | 18 | 18 | 18 | 18 | 15 | 18 | 18 |
| 32 | 15 | 15 | 15 | 18 | 18 | 18 | 18 | 18 | 18 | 15 | 18 | 18 |
| 33 | 15 | 15 | 15 | 18 | 18 | 18 | 18 | 18 | 18 | 18 | 18 | 18 |
| 34 | 15 | 15 | 15 | 18 | 18 | 18 | 18 | 18 | 18 | 18 | 18 | 18 |
| 35 | 15 | 15 | 15 | 18 | 18/15 | 18/15 | 18 | 18 | 18 | 18 | 18 | 18/15 |

**Supplementary Table S7:** Initial offset table after application of Integer Programming algorithm to a Pooled dataset in mESCs consisting of all genes. Offset table for application of Integer Programming algorithm to a Pooled dataset of *E. coli.*

| Fragment size | mESCs Pooled dataset | | | | *E. coli* Pooled dataset | | |
|---|---|---|---|---|---|---|---|
| | Frame 0 | Frame 1 | Frame 2 | | Frame 0 | Frame 1 | Frame 2 |
| **20** | 6/3 | 6/9 | 0/15 | | 12/9 | 15/0 | 9/0 |
| **21** | 6/0 | 6/0 | 0/18 | | 12/9 | 3/12 | 9/0 |
| **22** | 0/9 | 0/3 | 9/15 | | 12/0 | 12/0 | 9/0 |
| **23** | 6/15 | 9/0 | 18/9 | | 12/3 | 0/12 | 9/0 |
| **24** | 9/15 | 3/0 | 18/0 | | 12/3 | 12/0 | 0/9 |
| **25** | 9/6 | 9/6 | 12/15 | | 12/3 | 12/3 | 9/6 |
| **26** | 12/6 | 9/12 | 15/0 | | 12/3 | 3/0 | 9/3 |
| **27** | 12/9 | 12/21 | 15/9 | | 3/12 | 3/6 | 9/3 |
| **28** | 12/15 | 12/21 | 15/12 | | 3/12 | 3/6 | 3/0 |
| **29** | 15/12 | 15/12 | 18/15 | | 3/12 | 3/6 | 3/9 |
| **30** | **15** | 15/18 | 18/15 | | 12/9 | 9/3 | 9/6 |
| **31** | 15/12 | 15/18 | 18/15 | | 12/3 | 9/6 | 9/3 |
| **32** | 12/15 | 18/15 | **18** | | 12/9 | 3/9 | 9/3 |
| **33** | 15/12 | 18/15 | **18** | | 12/9 | 9/3 | 9/3 |
| **34** | 15/18 | 15/18 | 18/12 | | 12/9 | 9/12 | 9/6 |
| **35** | ND | ND | ND | | 12/9 | 12/9 | 9/6 |
| **36** | ND | ND | ND | | 12/15 | 9/12 | 9/12 |
| **37** | ND | ND | ND | | 12/6 | 12/9 | 9/12 |
| **38** | ND | ND | ND | | 12/15 | 12/15 | 9/12 |
| **39** | ND | ND | ND | | 12/15 | 12/3 | 9/3 |
| **40** | ND | ND | ND | | 12/15 | 15/3 | 9/3 |

*ND = Not Defined. The number of genes for certain $S$ and $F$ combinations are less than 10 and hence Integer Programming algorithm is not applied to these combinations.

**Supplementary Table S8:** A-site locations (nucleotide offsets from 5′ end) determined by applying the Integer Programming algorithm to the Pooled dataset in mESCs are shown as a function of fragment size and frame. The dataset consists of only genes that have a single isoform and only one translation start site. The top two offset values are listed for those $S$ and $F$ combinations in which the A-site location could not be uniquely determined. The description of the sub columns is the same as Supplementary Table S5.

<table>
<tr><th>Fragment Size</th><th colspan="3">Frame 0</th><th colspan="3">Frame 1</th><th colspan="3">Frame 2</th></tr>
<tr><td rowspan="2">28</td><td colspan="3" rowspan="2">15/12</td><td colspan="3" rowspan="2">15/12</td><td rowspan="2">15</td><td>S</td><td>R</td></tr>
<tr><td colspan="2">1 of 2</td></tr>
<tr><td rowspan="2">29</td><td rowspan="2">15</td><td>R</td><td>R</td><td colspan="3" rowspan="2">15/18</td><td colspan="3" rowspan="2">15/18</td></tr>
<tr><td colspan="2">2 of 2</td></tr>
<tr><td rowspan="2">30</td><td rowspan="2">15</td><td>R</td><td>R</td><td colspan="3" rowspan="2">15/18</td><td colspan="3" rowspan="2">18/15</td></tr>
<tr><td colspan="2">2 of 2</td></tr>
<tr><td rowspan="2">31</td><td rowspan="2">15</td><td>R</td><td>R</td><td rowspan="2">15</td><td>S</td><td>S</td><td rowspan="2">18</td><td>R</td><td>R</td></tr>
<tr><td colspan="2">2 of 2</td><td colspan="2">1 of 2</td><td colspan="2">2 of 2</td></tr>
<tr><td rowspan="2">32</td><td rowspan="2">15</td><td>R</td><td>R</td><td colspan="3" rowspan="2">18/15</td><td rowspan="2">18</td><td>R</td><td>R</td></tr>
<tr><td colspan="2">2 of 2</td><td colspan="2">2 of 2</td></tr>
<tr><td rowspan="2">33</td><td rowspan="2">15</td><td>R</td><td>R</td><td rowspan="2">18</td><td>R</td><td>R</td><td rowspan="2">18</td><td>R</td><td>R</td></tr>
<tr><td colspan="2">2 of 2</td><td colspan="2">2 of 2</td><td colspan="2">2 of 2</td></tr>
<tr><td rowspan="2">34</td><td colspan="3" rowspan="2">15/12</td><td rowspan="2">18</td><td>S</td><td>S</td><td rowspan="2">18</td><td>R</td><td>R</td></tr>
<tr><td colspan="2">2 of 2</td><td colspan="2">2 of 2</td></tr>
</table>

**Supplementary Table S9:** A-site offsets determined using the publicly available R packages – Plastid [4], RiboProfiling [5] and riboWaltz [6]. These methods generate a P-site offset as output for each fragment length. The A-site offsets below are obtained after adding 3nt to the P-site offsets.

| | ***S. cerevisiae* Pop data** | | | | ***S. cerevisiae* Pooled data** | | | | **mESCs Pooled data** | | |
|---|---|---|---|---|---|---|---|---|---|---|---|
| Fragment size | Plastid | RiboProfiling | riboWaltz | | Plastid | RiboProfiling | riboWaltz | | Plastid | RiboProfiling | riboWaltz |
| 20 | 16 | 7 | 16 | | 16 | 7 | 16 | | NA | NA | NA |
| 21 | 16 | 7 | 16 | | 16 | 7 | 13 | | NA | NA | NA |
| 22 | 16 | 7 | 15 | | 16 | 7 | 15 | | NA | NA | NA |
| 23 | 16 | 10 | 15 | | 16 | 10 | 15 | | NA | NA | NA |
| 24 | 16 | 10 | 17 | | 16 | 10 | 15 | | NA | NA | NA |
| 25 | 16 | 11 | 15 | | 16 | 11 | 15 | | 16 | 13 | 15 |
| 26 | 16 | 11 | 16 | | 16 | 11 | 16 | | 16 | 14 | 16 |
| 27 | 16 | 14 | 14 | | 16 | 14 | 14 | | 16 | 15 | 15 |
| 28 | 16 | 15 | 15 | | 16 | 15 | 15 | | 13 | 16 | 15 |
| 29 | 16 | 16 | 16 | | 16 | 16 | 16 | | 13 | 6 | 15 |
| 30 | 16 | 16 | 16 | | 16 | 16 | 16 | | 15 | 15 | 15 |
| 31 | 16 | 16 | 16 | | 16 | 16 | 16 | | 13 | 13 | 16 |
| 32 | 16 | 17 | 17 | | 16 | 17 | 17 | | 16 | 13 | 16 |
| 33 | 16 | 17 | 17 | | 16 | 17 | 17 | | 16 | 14 | 16 |
| 34 | 16 | 13 | 15 | | 16 | 13 | 15 | | 17 | 13 | 14 |
| 35 | 16 | 13 | 16 | | 16 | 13 | 15 | | NA | NA | NA |

**Supplementary Table S10:** Median normalized ribosome densities for 61 codon types were correlated with tRNA abundance for the Integer Programming method and 9 other contemporary methods. The tRNA abundance values were obtained from Table S2 of study of Weinberg and co-workers [7].

| Method | Spearman's rho | p-value |
|---|---|---|
| Integer Programming | -0.583 | $6.39 \times 10^{-5}$ |
| Heuristic +18 | -0.581 | $6.76 \times 10^{-5}$ |
| Plastid | -0.580 | $6.98 \times 10^{-5}$ |
| RiboProfiling | -0.575 | $8.55 \times 10^{-5}$ |
| riboWaltz | -0.574 | $8.65 \times 10^{-5}$ |
| Hussmann | -0.571 | $9.53 \times 10^{-5}$ |
| Martens | -0.571 | $9.82 \times 10^{-5}$ |
| Heuristic +15 | -0.570 | $9.94 \times 10^{-5}$ |
| ribodeblur | -0.570 | $9.94 \times 10^{-5}$ |
| Scikit-ribo | -0.567 | $1.09 \times 10^{-4}$ |
| Rpbp | -0.566 | $1.12 \times 10^{-4}$ |
| Center-weighted | -0.517 | $5.31 \times 10^{-4}$ |
